# A solution to temporal credit assignment using cell-type-specific modulatory signals

**DOI:** 10.1101/2020.11.22.393504

**Authors:** Yuhan Helena Liu, Stephen Smith, Stefan Mihalas, Eric Shea-Brown, Uygar Sümbül

## Abstract

Animals learn and form memories by jointly adjusting the efficacy of their synapses. How they efficiently solve the underlying temporal credit assignment problem remains elusive. Here, we re-analyze the mathematical basis of gradient descent learning in recurrent spiking neural networks (RSNNs) in light of the recent single-cell transcriptomic evidence for cell-type-specific local neuropeptide signaling in the cortex. Our normative theory posits an important role for the notion of neuronal cell types and local diffusive communication by enabling biologically plausible and efficient weight update. While obeying fundamental biological constraints, including separating excitatory vs inhibitory cell types and observing connection sparsity, we trained RSNNs for temporal credit assignment tasks spanning seconds and observed that the inclusion of local modulatory signaling improved learning efficiency. Our learning rule puts forth a novel form of interaction between modulatory signals and synaptic transmission. Moreover, it suggests a computationally efficient learning method for bio-inspired artificial intelligence.

## Introduction

Animals learn and form memories by adjusting the efficacy of particular subsets of the myriad synaptic connections that establish their nervous system architectures. Borrowing terminology that was introduced in the early days of artificial intelligence [1], identification of the connection subset necessary for adaptive learning has come to be known by neuroscientists, too, as “credit assignment” – that is, the assignment of credit (or blame) to particular synaptic connections as needed to guide the strengthening (or weakening) to achieve adaptive memory formation. Unfortunately, the mechanisms of credit assignment for biological networks remain deeply enigmatic [2, 3, 4].

Mathematical “gradient backpropagation” algorithms [5, 6] now solve the problem of credit assignment for artificial neural networks sufficiently well to have ushered in an era of shockingly powerful artificial intelligence, but the training of networks by these algorithms still face oppressive scalability issues. Computer scientists therefore continue to look to neuroscience for inspiration regarding new approaches to credit assignment. Neuroscientists meanwhile still struggle to see how the computer scientists’ backpropagation approach to credit assignment could be implemented by the brain’s “hardware”. Inspired by new findings regarding neuromodulatory signaling from single-cell RNA sequencing analysis of mouse brain gene expression [7, 8], we have developed and propose here a simple theory which may contribute to a new wave of progress in understanding biological credit assignment and may also serve to inspire more efficient credit assignment in the artificial intelligence realm.

One conspicuous gap between computational models of neuronal networks and experimental data appears in the concept of cell types. Recently, remarkable diversity and stereotypy have been observed in neuronal phenotypes [9,10,11,12, 13]. Despite attempts at bridging this gap by explicitly studying discrete cell types as part of the computational model [14], an overarching role for cell types in synaptic plasticity is not yet known.

Hebbian plasticity, a local learning rule that prescribes lasting change in synaptic strength based on correlations of spike timing between particular presynaptic and postsynaptic neurons, has long been recognized as one biologically plausible basis for assigning credit to particular synapses during memory formation [15, 16]. Both experimental and theoretical investigations now indicate forcefully, however, that a Hebbian rule alone is insufficient: impacts of correlated spike timing must be augmented by one or more additional, modulatory factors [17,18,19,20,2,21,22,23,24,25,26,27]. Dopamine, a neuromodulatory monoamine secreted by axons ramifying widely from midbrain in response to salient “rewarding” or “surprising” events, has emerged as the most prominent candidate for such a factor [28]. Neurons respond to secreted dopamine strictly via activation of G protein-coupled receptors (GPCRs), which act via intracellular messengers upon ion channels, secretory machinery and gene expression over widely varied time scales to regulate membrane excitability, synaptic strength and the temporal parameters of Hebbian plasticity rules.

Five different genes encoding dopamine-selective GPCRs are expressed in highly differential, cell-type-specific fashion by central nervous system (CNS) neurons, but so are hundreds of other genes encoding GPCRs of widely varying ligand selectivity [29]. A majority of these other GPCRs are selective for diffusible ligands that are secreted in response to neuronal activity and many are recognized, like dopamine, as potent neuromodulators [30, 31, 32, 33]. These neuromodulatory GPCRs include receptors selective for other monoamines such as serotonin, norepinephrine and histamine, metabotropic receptors for the neurotransmitters glutamate and GABA, the muscarinic acetylcholine receptors, endocannabinoid receptors, and receptors selective for neuropeptides [34]. Most of these other GPCRs have downstream signaling actions quite similar to those of the dopamine-selective GPCRs and many furthermore have been shown to impact memory across a wide range of animal species [35, 36, 37]. Thus, numerous neuromodulatory GPCRs beyond the dopamine receptors merit serious consideration as potential additional factors contributing to credit assignment. Only a few are likely to act like dopamine as top-down reward signals, but many are likely to distribute information about local network activity locally in ways that might contribute significantly to the speed and efficiency of credit assignment.

The present work was inspired especially by new single-cell RNA sequencing (scRNA-seq) evidence for surprisingly abundant and diverse expression of neuropeptide precursor protein (NPP) and NP-selective GPCR (NP-GPCR) genes in virtually all mouse neocortical neurons [7, 8]. Active neuropeptide (NP) molecules are proteolytic fragments of proteins encoded by NPP genes and released from neurons in activity-dependent fashion. NPs comprise the largest family of diffusible ligands for neuromodulatory GPCRs [38, 39] and many experimental findings have implicated them in memory storage processes [37, 40, 41]. The fundamental importance of NP signaling is supported by their extremely deep conservation across all animal species [42] and by extremely high levels of NPP-encoding mRNA abundance in individual neurons [7]. The most widely known NPs are the opioids (enkephalins, endorphins and others) and the pro-social peptide oxytocin, but there are many others [43]. In mammalian CNS, products of approximately 50 NPP genes act via NP-GPCR proteins encoded by approximately 80 different NP-GPCR genes. The scRNA-seq data further show that almost all mouse CNS neurons express several NPPs and several NP-GPCRs and that, for both families, the combinations expressed are extraordinarily cell-type-specific [7]. Though secreted NP molecules must diffuse far enough through brain interstitial spaces to reach many individual synapses connecting neurons representing multiple types [44, 45], tightly defined patterns of sparse, cell-type-specific expression of particular signal source (NPP) and signal receiver (NP-GPCR) genes nonetheless arguably define directed graphs of connectivity between nearby neurons of particular types. These findings thus suggest a “multidigraph” view of cortical computation and plasticity, founded upon interplay between fast synaptic and slow neuromodulatory networks, definable by multiple directed graphs sharing cell types as common nodes [8] (Figure 1). Here we explore this prospect theoretically in the hope of motivating and facilitating experimentation and developing experimentally testable predictions.

**Figure 1:**
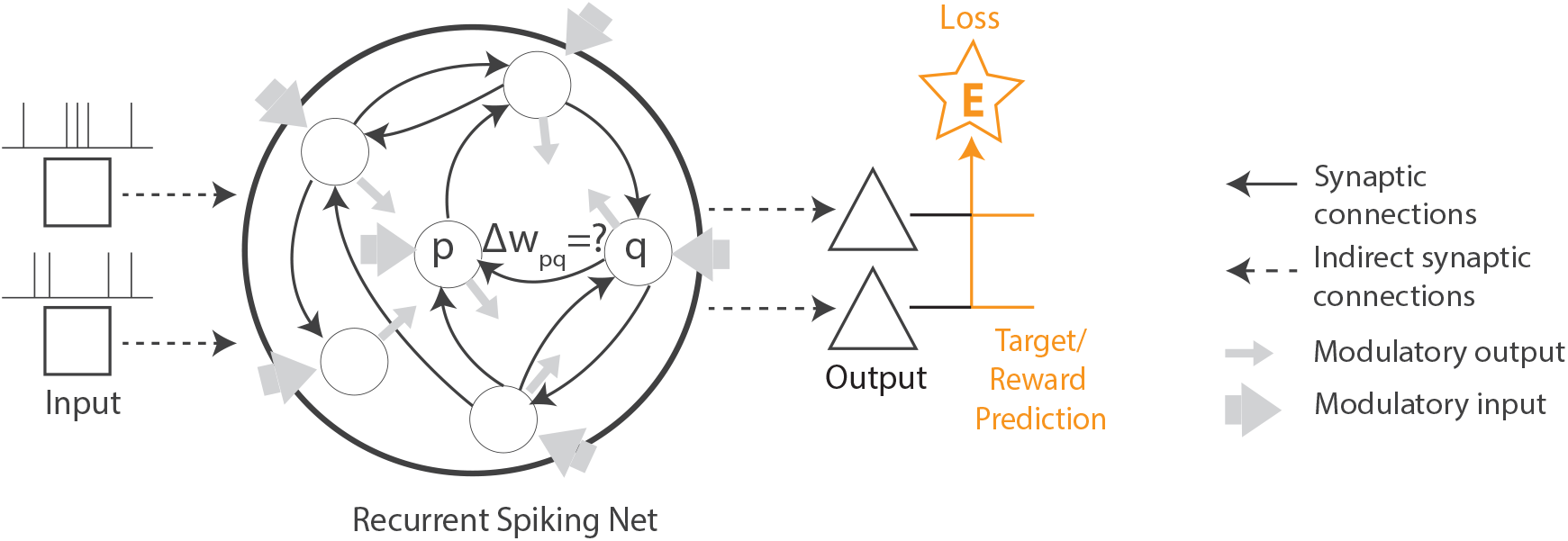
Temporal credit assignment through the interplay of synaptic transmission and modulatory signaling. The network is tasked with producing a desired output signal given a certain input. The challenge involves determining how much each weight (out of potentially thousands or millions of connections) is responsible for overall network performance, so as to guide the strengthening and weakening of the individual weights. In this view of the neuronal network as a stack of synaptic and modulatory networks, learning (i.e the update of synaptic weights *w*) is shaped by both local synaptic activity and modulatory signaling.

A major step forward in biologically plausible learning in recurrent neural networks (RNNs) – widely-adopted high dimensional dynamical models of neural circuits for robust performance in temporal tasks [46] – was brought by two recent theoretical studies that developed local and causal learning rules [47, 48]. These studies derived local approximations to gradient-based learning in RNNs by requiring synaptic weight updates to depend only on local information about pre- and postsynaptic activities in addition to a top-down learning signal pertaining to network output error. While Murray approximated RTRL for rate-based networks [47], Bellec *et al*. approximated BPTT to train recurrent spiking neural networks (RSNNs) [48]; this spike-based communication yields greater biological plausibility and energy efficiency [49,50,51,52,53].

Building on these recent advances, we test the plausibility of the abstract multidigraph concept by formulating it into an explicit computational model and describe a computational role for cell types in synaptic learning as part of our model. More specifically, we truncate the RTRL algorithm to remove nonlocal dependencies, but include modulatory terms respecting neuronal types to provide nonlocal information in the form of diffusive signaling. (truncations of the RTRL algorithm have received recent attention from the machine learning and neuromorphic hardware communities [54, 55].) Our multidigraph learning rule (MDGL) generalizes multi-factor learning, in which a Hebbian component is combined with local cell-type-specific modulatory signals in addition to the top-down instructive signals. We train the multiple-cell-type RSNNs with MDGL to perform tasks involving temporal credit assignment over a timescale of seconds. Although we focus on supervised learning, our theory can be extended to reinforcement learning settings [48]. Our proof-of-concept implementation of MDGL shows significant improvements over previous literature and advances the field of biologically plausible temporal credit assignment. From a neuroscience perspective, our study proposes a new model of cortical learning shaped by the interplay of local modulatory signaling and synaptic transmission, and potentially brings us closer to understanding biological intelligence. From a computer science perspective, our method offers an energy efficient method for on-chip neuro-inspired AI.

## Results

### Mathematical basis of multidigraph learning in RSNNs

To study the basic principles governing plasticity in neuronal circuits, we use a simple and widely adopted recurrent neural network model with the addition that we endow neurons with local modulatory signaling and cell type-specific cognate reception (Figure 1). An overview of our biologically plausible learning framework along with other methods investigated in this work are summarized in Figure 2.

**Figure 2:**
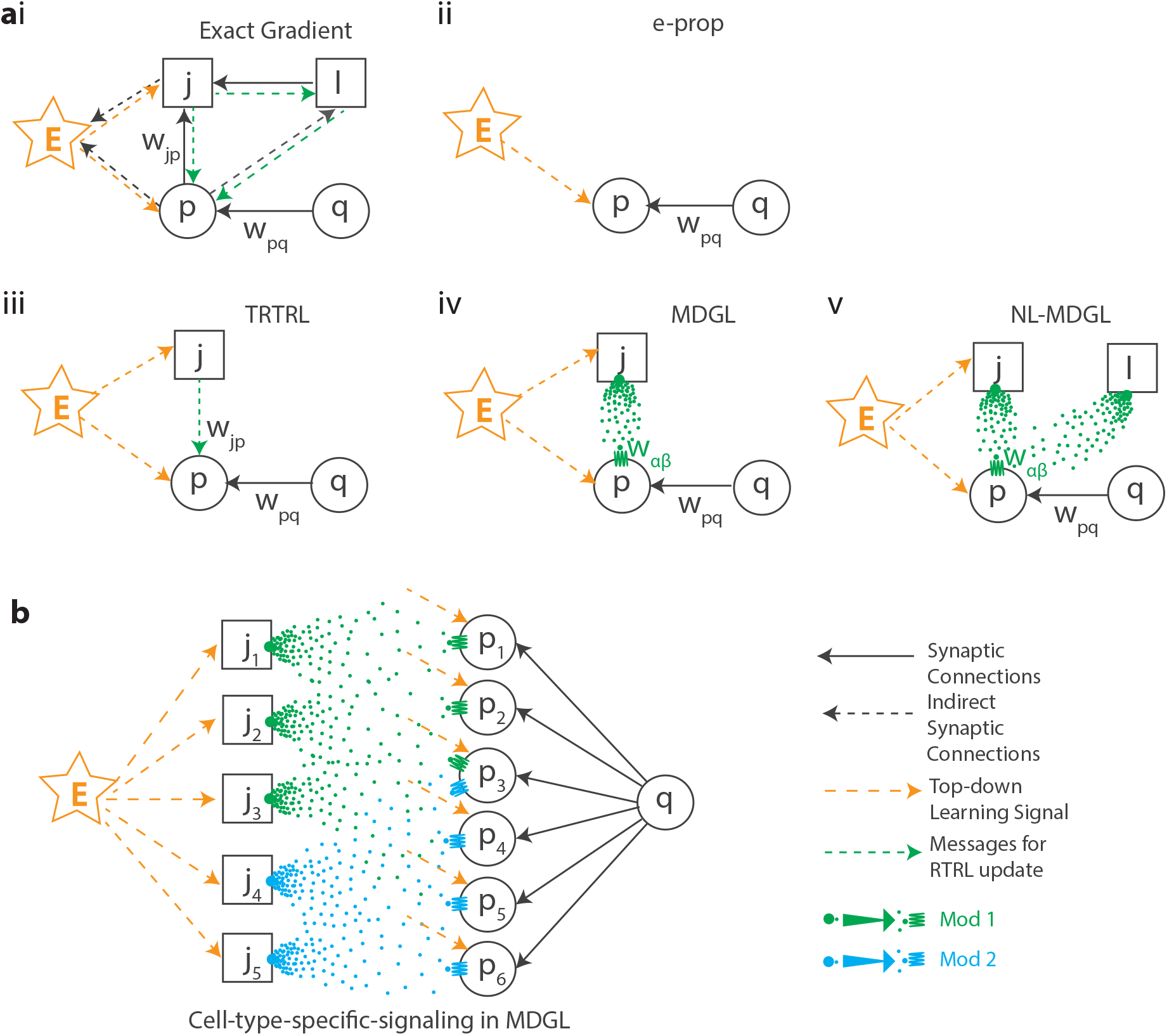
Biologically plausible temporal credit assignment using cell-type-specific modulatory signals. a) Learning rules investigated include (i) the exact gradient: updating weight *w_pq_*, the synaptic connection strength from presynaptic neuron *q* to postsynaptic neuron *p*, involves nonlocal information inaccessible to neural circuits, i.e. the knowledge of activity for all neurons *j* and *l* in the network. This is because *w_pq_* affects the activities of many other cells through indirect connections, which will then affect the network output at subsequent time steps. As in Figure 1, *E* quantifies the network’s performance. (ii) E-prop, a state-of-the-art local learning rule, restricts weight update to depend only on pre- and post-synaptic activity as well as a top-down learning signal. (iii) Our truncated weight update (TRTRL) includes dependencies within one connection step, which are omitted in e-prop. (iv) Our multi-digraph learning rule (MDGL) incorporates local cell-type-specific modulatory signaling through which the activity of neuron j can be delivered to neuron p. (v) NL-MDGL: A nonlocal version of MDGL, where modulatory signal diffuses to all cells in the network. b) Illustration of cell-type-specific modulatory signaling in MDGL: source neurons *j*_1_, *j*_2_ and *j*_3_ express the same precursor type, and likewise for *j*_4_ and *j*_5_. Neurons *p*_1_ to *p*_6_ can be grouped into three types based on the different combination of receptors they express. This results in a modulatory network described by a cell-type-specific channel gain on top of the classical synaptic network.

The RSNN used in this study (Figures 1 and 3a) takes *N*_*in*_ spiking inputs *x*_*i,t*_ for *i* = 1, 2, …*N*_*in*_ and *t* = 1, 2, …, *T*, where *x*_*i,t*_ assumes the value 1 if input unit *i* fires at time *t* and 0 otherwise. These inputs are sent to the spiking recurrent units. Internal to recurrent unit *j* is state *s*_*j,t*_, which denotes the internal state (e.g. membrane potential for LIF neurons) of cell *j* at time *t*. The output of the *j*^th^ recurrent neuron at time *t, z*_*j,t*_ for *j* = 1, 2, …*N*, also takes the value 1 if recurrent neuron *j* fires at time *t* and 0 otherwise. The recurrent activity is read out to graded output *y*_*k,t*_ for *k* = 1, 2, …*N*_*y*_, whose performance for a given task incurs a feedback signal *E* (e.g., error or negative reward). Throughout, many variables of interest display temporal dependencies and spatial (e.g., cell index *p*, synapse index *pq* for connections from presynaptic neuron *q* to postsynaptic neuron *p*) dependencies. Equations governing the dynamics of the non-adaptive and adaptive leaky integrate-and fire (LIF) neurons used in this paper are given in Methods.

**Figure 3:**
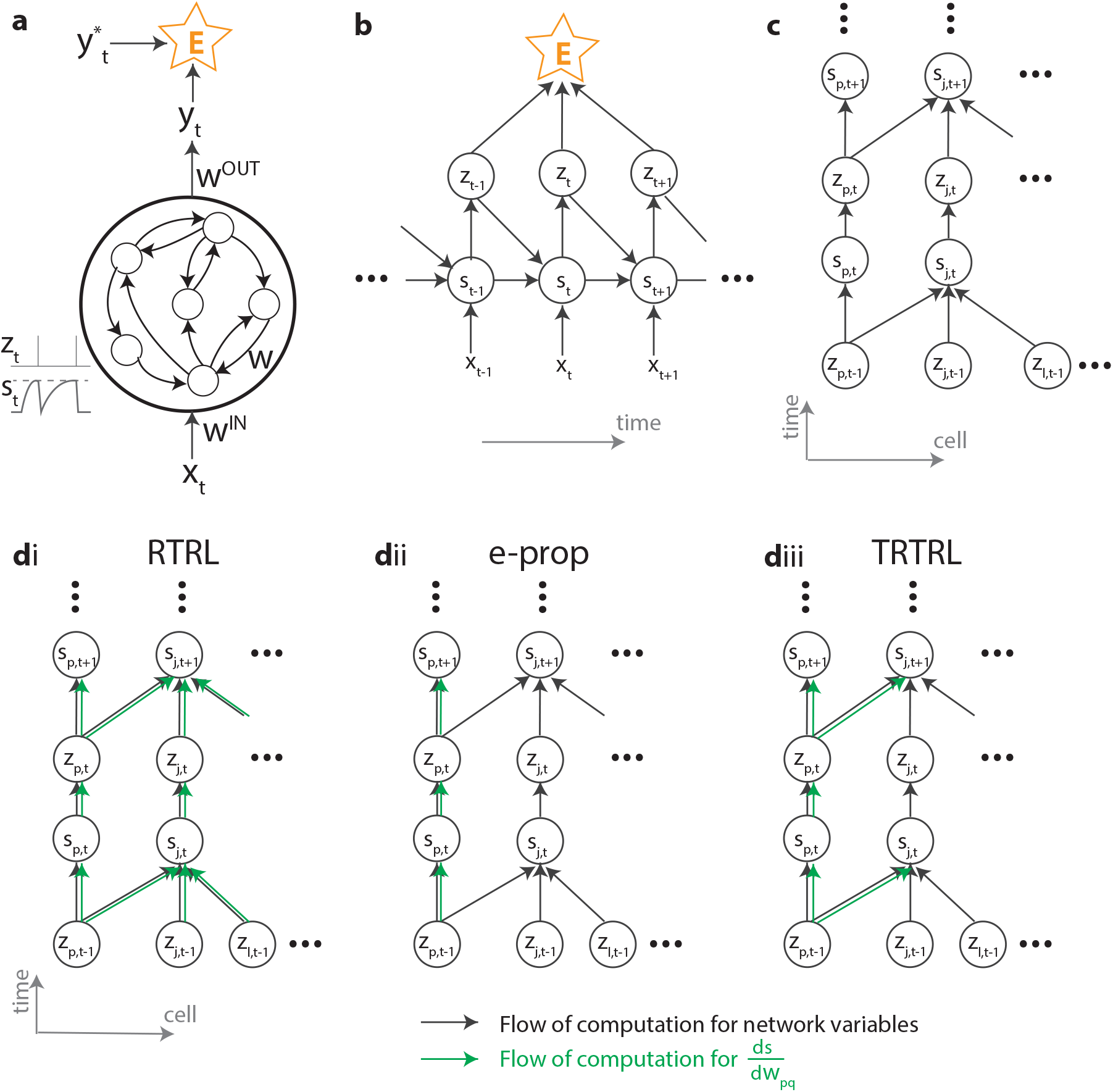
Computational graph and gradient propagation. a) Schematic illustration of the recurrent neural network used in this study. b) The mathematical dependencies of input *x*, state *s*, neuron spikes *z* and loss function *E* unwrapped across time. c) The dependencies of state *s* and neuron spikes *z* unwrapped across time and cells. d) The computational flow of d *s/* d *w*_*pq*_ is illustrated for (di) exact gradients computed using RTRL, (dii) e-prop and (diii) our truncation in Eq. 2, where dependency within one connection step has been kept. Black arrows denote the computational flow of network states, output and the loss; for instance, the forward arrows from *z*_*t*_ and *s*_*t*_ going to *s*_*t*+1_ are due to the neuronal dynamics equation in Eq. M1. Green arrows denote the computational flow of d *s/* d *w*_*pq*_ for various learning rules.

We study iterative adjustment of all synaptic weights (input weights *w*^*IN*^, recurrent weights *w* and output weights *w*^*OUT*^) using gradient descent on *E*, which involves a weight update in the direction of error gradient (see Methods for gradient descent and detailed definitions of *E*). This error gradient can be calculated with BPTT or RTRL by unwrapping the RSNN dynamics over time; this unwrapping is needed because weights influence past network activity, which then influences present and future activity through Eq. M1, Methods (Figure 3b). While these two algorithms yield equivalent results, their bookkeeping for the gradient calculations differs [47]. Gradient calculations in BPTT depend on future activity, which poses an obstacle for online learning and biological plausibility. Unlike BPTT, the computational dependency graph of RTRL is causal. Therefore, we focus our analysis on RTRL and factor the error gradient across time and space (Eqs. M8 and M9, Methods).

Key problems that RTRL poses to biological plausibility and computational cost reside in the factor 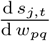 that arises during the factorization of the gradient (Eq. M8 and Eq. M9). The factor 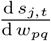 keeps track of all direct and indirect dependencies of neuron state *j* on weight *w*_*pq*_. In other words, this factor accounts for both the spatial and temporal dependencies in RSNNs: state dependencies across time *t*, as explained above, result from unwrapping the temporal dependencies illustrated in Figure 3b; state dependencies across space, however, are due to the indirect dependencies (of all *z*_*t*_ on *w* and all *z*_*t*_*’* (*t*^*’*^ < *t*)) arising from recurrent connections (Figure 3c). These recurrent dependencies are all accounted for in the 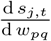 factor, which can be obtained recursively as follows:

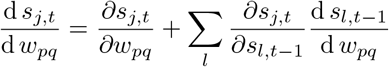

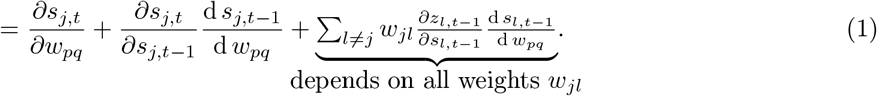

Thus, the factor 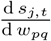 is a memory trace of all inter-cellular dependencies (Figures 3di, 2ai), requires *O*(*N* ^3^) memory and *O*(*N* ^4^) computations. This makes RTRL expensive to implement for large networks. Moreover, this last factor poses a serious problem for biological plausibility: it involves nonlocal terms, so that knowledge of all other weights in the network is required in order to update the weight *w*_*pq*_.

To address this, Murray [47] and Bellec *et al*. [48] (“e-prop”) dropped the nonlocal terms so that the updates to weight *w*_*pq*_ would only depend on pre- and post-synaptic activity (Figures 3dii, 2aii), and applied this truncation to train rate-based and spiking neural networks, respectively. While both works succeed in improving over previous biologically plausible learning rules, a significant performance gap with respect to the full BPTT/RTRL algorithms remains. This gap is not surprising given that both algorithms account only for pre- and post-synaptic activities, ignoring – by design – the many potential contributions to optimal credit assignment from neurons that do not participate directly in the synapse of interest.

### A potential role for cell type-specific modulatory signals

To reveal a potential role for cell-type-based modulatory signals in synaptic plasticity, we begin by partially restoring non-local dependencies between cells – those within one connection step. This is the *truncated* RTRL framework (Figures 3diii, 2aiii), and the memory trace term 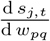 becomes

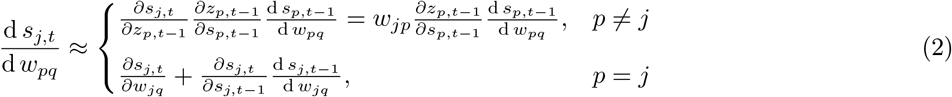

Thus, when *j* = *p*, our truncation implements 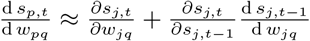, which coincides with e-prop. Eq. 2 adds the case when *p* ≠ *j*, for which 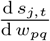 was simply set to 0 in e-prop. We note that the truncation in Eq. 2 resembles the n-step RTRL approximation recently proposed in [54], known as SnAP-n, which stores 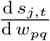 only for *j* such that parameter *w*_*pq*_ influences the activity of unit *j* within *n* time steps. The computations of SnAp-n converge to those of RTRL as n increases. Our truncation in Eq. 2 is similar to SnAp-n with *n* = 2 with two differences: (i) we apply it to spiking neural networks, (ii) we drop the previous time step’s Jacobian term 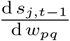, which would necessitate the maintenance of a rank-three (“3-d”) tensor with costly storage demands (*O*(*N* ^3^)) and for which no known biological mechanisms exist. Thus, the truncation in Eq. 2 requires the maintenance of only a rank-two (“2-d”) tensor specific to synapse *pq*, which can be realized via an eligibility trace as we explain next.

By substituting equation Eq. 2 into Eq. M8 and Eq. M9, we approximate the overall gradient as

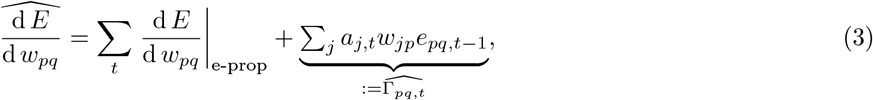

where *a*_*j,t*_ (Eq. M12) denotes the activity-dependent modulatory signal emitted by neuron *j* at time *t* and *e*_*pq,t*_ (Eq. M13) is the *eligibility trace* maintained by postsynaptic cell *p* to keep a memory of the preceding activity of presynaptic cell *q* and postsynaptic cell *p* (Methods). There have been numerous suggestions that some persistent “eligibility trace” must exist to bridge the temporal gap between correlated firing at specific synapses and later-arriving, dopamine-like reward signals [21, 26, 56,57,58,59]. In Eq. 3, the first term alone gives exactly the e-prop synaptic update rule. The second term, which we define as 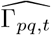 is a synaptically non-local term that is ignored by e-prop. As seen in Eq. 3, our truncation requires maintaining a {*p, q*} -dependent double tensor (for *e*_*pq,t*_) instead of a triple one, thereby reducing the memory cost of RTRL from *O*(*N* ^3^) to *O*(*N* ^2^).

Importantly, we observe that, for the update to synapse *w*_*pq*_ in Eq. 3, the terms that depend on cells *j only appear under a sum*. Therefore, the mechanism updating the synapse (*pq*) does not need to know the individual terms indexed by *j*. Rather, only their sum suffices. This observation is key in hypothesizing a role for diffuse neuromodulatory signalling as an additional factor in synaptic plasticity. As discussed above, multiple different neuromodulators diffuse in the shared intercellular space [7, 8]; this provides a natural biological substrate for the transmission of weighted, summed signals.

While it is tempting to consider the first factors in 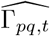, *a*_*j,t*_*w*_*jp*_, as the modulatory signal emitted by neuron *j*, the involvement of the synapse from neuron *p* via *w*_*jp*_ and a lack of known mechanisms in calculating this neuron-specific composite signal suggest that this is unlikely to be a biological solution. Instead, inspired by the cell-type-specific (rather than neuron-specific) affinities for peptidergic neuromodulation [7, 8], we propose to approximate the signaling gain *w*_*jp*_ in Eq. 3 by the average value *w*_*αβ*_ across its pre- and postsynaptic cell types. More specifically, when postsynaptic cell *j* belongs to type *α* and presynaptic cell *p* belongs to type *β*, we approximate neuron-specific weight *w*_*jp*_ with cell-type-specific gain *w*_*αβ*_ =< *w*_*jp*_ >_*j∈α,p∈β*_. We hypothesize that *w*_*αβ*_ represents the affinity of the G-protein coupled receptors expressed by cells of type *β* to the peptides secreted by cells of type *α* (Figure 2aiv, b). Finally, the local diffusion hypothesis discussed in [39] suggests a further approximation, in which this type of signaling is registered only by local synaptic partners and therefore preserves the connectivity structure of *w*_*jp*_:

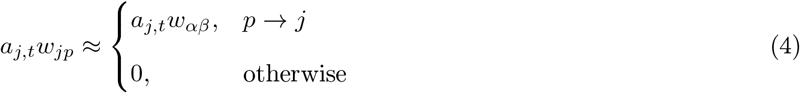

where *p* →*j* denotes that there is a synaptic connection from neuron p to j. In other words, while *a*_*j,t*_ is emitted by neuron *j*, its effect on neuron *p* through diffusion, as part of a sum, is given by Equation 4.

Bringing these together, the gradient estimate at time *t* due to our learning rule is given as

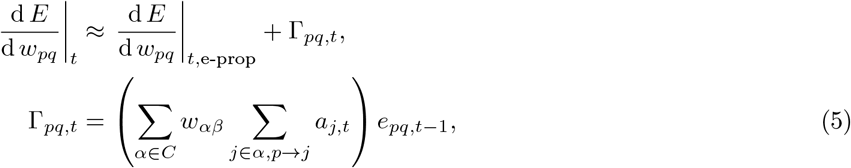

where neuron *p* is of type *β, C* denotes the set of neuronal cell types, Γ_*pq,t*_ approximates the second term in Eq. 3 with cell-type-specific weight averages. Thus, our update rule suggests a new additive term to compute the plasticity update at synapse *pq* at time *t*, Γ_*pq,t*_, which calculates multiplicative contributions of the modulatory signal *a*_*j,t*_ secreted by neuron *j*, the affinity of receptors of cell type *β* to ligands of type *α, w*_*αβ*_, and the eligibility trace at the synapse *pq, e*_*pq,t*_. While Eq. 5 suggests an “online” implementation with the update at *t*, the factor *a*_*j,t*_ cannot be calculated causally unless the output is not leaky. Supplementary Note 1 derives an online update for the more general case, which has the same form as Eq. 5.

In summary, we have proposed a new rule for updating a synapse *w*_*pq*_, which we refer to as the multidigraph learning rule, or MDGL, and represent by Δ*w*_*pq* | *MDGL*_. As illustrated in Fig. 2, this begins with the same basic structure as e-prop, which has the following form:

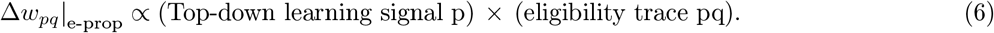

Δ *w*_*pq*_ |_MDGL_ then adds a term Γ_*pq*_, whose form matches that of diffusive communication via a cell-type specific modulatory network. Taking account of the fact that the activity-dependent modulatory signal emission *a*_*j,t*_ is a combination of the top-down learning signal received by neuron *j* and its activity (see Eq. M12 and detailed component breakdown in Supplementary Note 2):

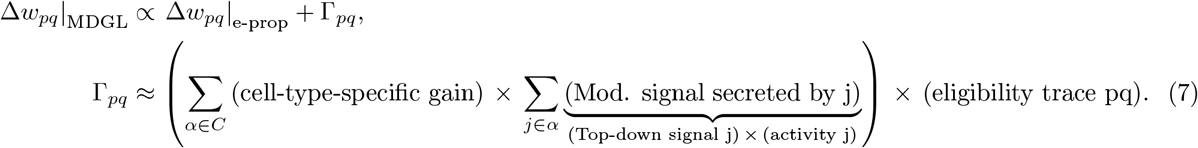

Thus, the Hebbian eligibility trace is not only compounded with top-down learning signals – as in modern biologically plausible learning rules [25] – but also integrated with cell-type-specific, diffuse modulatory signals. This creates a unified framework that integrates the eligibility trace, local and top-down modulatory signals into a new multi-factor learning rule.

### Simulation of multidigraph learning in RSNNs

To test the efficiency of the MDGL formulation for synaptic plasticity, we apply it to three well-known supervised learning tasks involving temporal processing: pattern generation, a delayed match to sample task, and evidence accumulation. We use two main cell classes, inhibitory (I) and excitatory (E) cells, and obey experimentally observed constraints: cells have synapses that are sign constrained with 80% of the population being excitatory and the rest inhibitory. We further endow a fraction of the E cells with threshold adaptation [48] to mimic the hierarchical structure of cell types that has been established empirically [9] through its simple example of two main cell types (E and I), one of which has two subtypes (E cells with and without threshold adaptation). Also, refractoriness and synaptic delay are incorporated into all cells’ dynamics. Moreover, overall connection probability in the RSNN is also constrained, reflecting the sparse connectivity in neuronal circuits [60, 61]. In the main text, all simulated tasks are constrained at 10% sparsity, and the sparsity parameter is varied in Figure S2 in the supplementary materials. This connection sparsity is maintained by fixing inactive synapses with 0 weights. Unlike the stochastic rewiring (Deep R) algorithm [62, 63], our implementation does not allow for rapid and random formation of new synapses after each experience, further increasing biological plausibility.

To study the impact of our learning rule on network performance and dissect the effects of its different components, we train RSNNs using five different approaches for each task (Figure 2): (i) BPTT, which updates weights using exact gradients; (ii) E-prop [48], the state-of-the-art method for biologically plausible training of RSNNs; (iii) TRTRL, the truncated RTRL given in Eq. 3 without the cell-type approximation; (iv) MDGL, which incorporates the cell type approximation given in Eq. 5 using only two cell types; (v) NL-MDGL, a nonlocal version of MDGL, where the gain is replaced by *w*_*αβ*_ =< *w*_*jp*_ > _*j ∈α,p∈β*_ even for *w*_*jp*_ = 0 so that the modulatory signal diffuses to all cells in the network. While the naive implementation of *a*_*j,t*_ depends on future errors in Figures 4–7, we derive an online approximation in Supplementary Note 1 and demonstrate that it does not lead to performance degradation (Figure S3).

**Figure 4:**
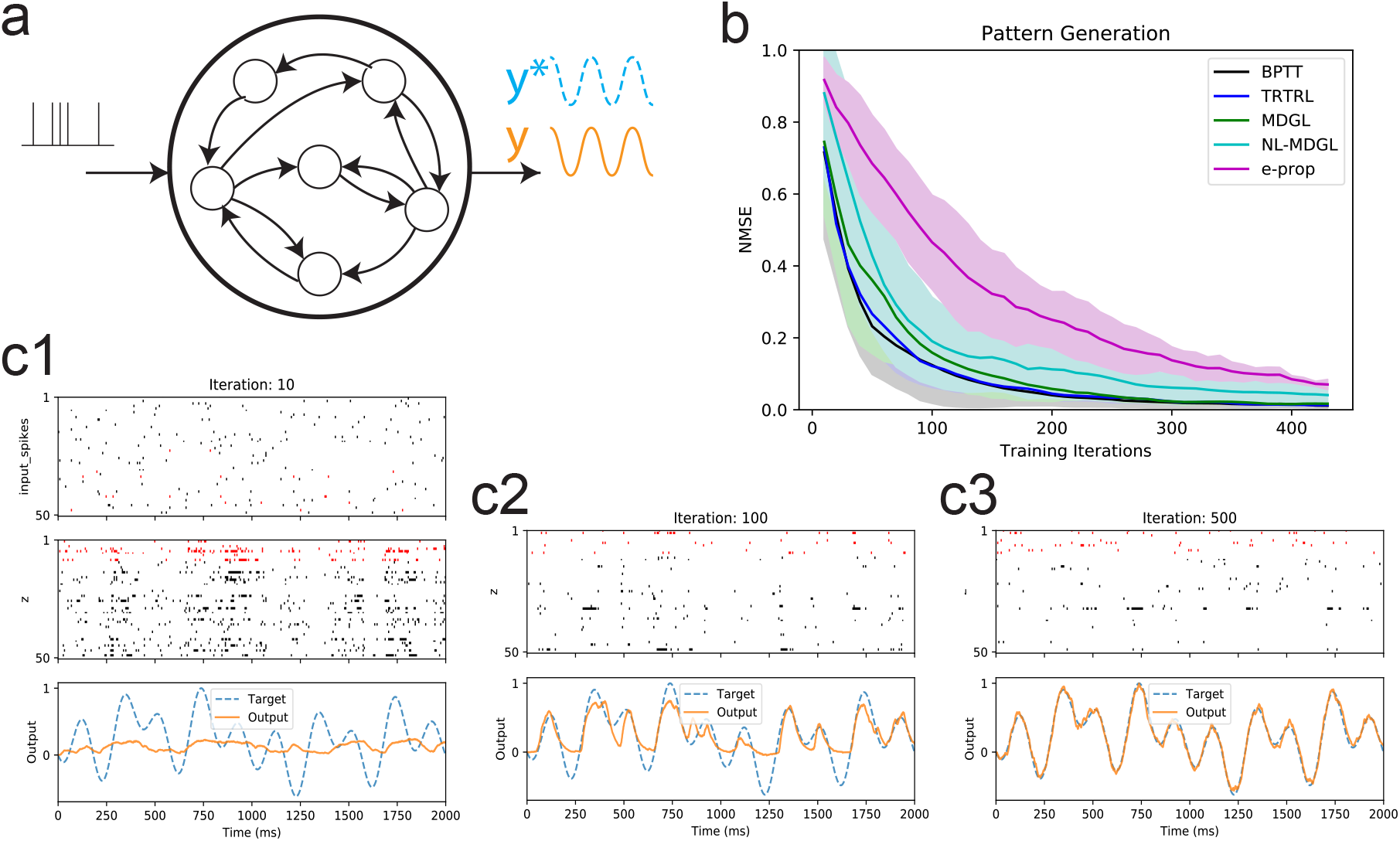
Pattern generation task. a) Task setup [65]: a network is trained to produce a target output pattern over time. The target is formed from a sum of five sinusoids. b) Normalized mean squared error (NMSE) over training iterations is illustrated for the five learning rules (Fig. 2). Solid lines show the mean and shaded regions show the standard deviation across five runs, with different target output, frozen Poisson input and weight initialization between runs (but fixed within each run). Comparing the performance of e-prop with the MDGL method suggests that the addition of cell-type-specific modulatory signals expedites the learning curve. c) Dynamics of the input, output and recurrent units are shown after 1, 100 and 500 iterations of training using the MDGL method. Raster plots are shown for 50 selected sample cells, and E cells and I cells are color coded using black and red, respectively. All recurrent units have fixed thresholds for this task. Recurrent unit spikes are irregular throughout training. Network output approaches the target as training progresses.

For each learning rule, we train the following network parameters: input, recurrent and output weights. All approaches update the output weights using backpropagation since the nonlocality problem in Eq. 1 applies only to the update of input and recurrent weights. (For updating the weights of a single output layer, random feedback alignment [64] has also been shown to be an effective and biologically plausible solution.)

### Multidigraph learning in RSNNs improves efficiency in a pattern generation task

We first trained a RSNN to produce a one-dimensional target output, generated from the sum of five sinusoids, given a fixed Poisson input realization. The task is inspired by that used in [65]. We compare the training for the five learning rules illustrated in Fig. 2 and described above.

While the target output and the random Poisson input realization is fixed across training iterations, we change these along with the initial weight for different training runs, and illustrate the learning curve mean and standard deviation in Figure 4b across five such runs. Here, the learning curve displays the normalized mean squared error (NMSE) between the actual and desired output over training iterations. We observe that our methods, TRTRL (blue) and MDGL (green), reduce NMSE faster over training iterations compared to e-prop (magenta). Also, removing the locality of the modulatory signal (NL-MDGL) degraded the efficiency, although learning with spatially non-specific modulation still outperformed that without the modulatory signal (e-prop).

We highlight the fact that approximating the nonlocal learning rule of TRTRL with diffuse, modulatory signaling among two cell types results in only a moderate degradation of performance. To better understand this observation, we conduct an analysis of the similarity between the 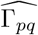 term computed by TRTRL in Eq. 3 and its cell-type-based approximation Γ_*pq*_ computed by MDGL. We quantify this via the alignment angle, which describes the similarity between the directions of the two vectors (Supplementary Note 3, Supplementary Table S1). The significant alignment between these two signals, despite the underlying vectors lying in very high dimensional spaces of synaptic weights, demonstrates that the two methods compute similar gradients, thus shedding light on the similarity between their learning curves.

Figure 4c illustrates the input, recurrent network activity, and target versus actual outputs trained using the MDGL method after training for 10, 100 and 500 iterations. Recurrent unit spikes appear to be reasonably irregular over time, broadly consistent with what is typically observed biologically [66, 67], with no obvious patterns of system-wise synchrony throughout training. Finally, the network output approaches the target as training progresses (lower panels).

### Multidigraph learning in RSNNs improves efficiency in a delayed match-to-sample task

To elucidate how neural networks with the learning rules at hand can be trained to integrate a history of past cues to generate responses that impact a reward delivered later, we considered a special case of the delayed match to sample task described in [68]. Here, two cue alternatives are encoded by the presence and absence of input spikes. Our implementation of the task began with a brief fixation period (no cues) followed by two sequential cues, each lasting 0.15s and separated by a 0.75s delay (Figure 5a). A cue of value 1 was represented by 40Hz Poisson spiking input, whereas a cue of value 0 was represented by the absence of input spiking. The network was trained to output 1 (resp. 0) when the two cues have matching (resp. non-matching) values. That is, the RSNN was trained to remember the first cue and learn to compare it with the second cue delivered at a later time.

**Figure 5:**
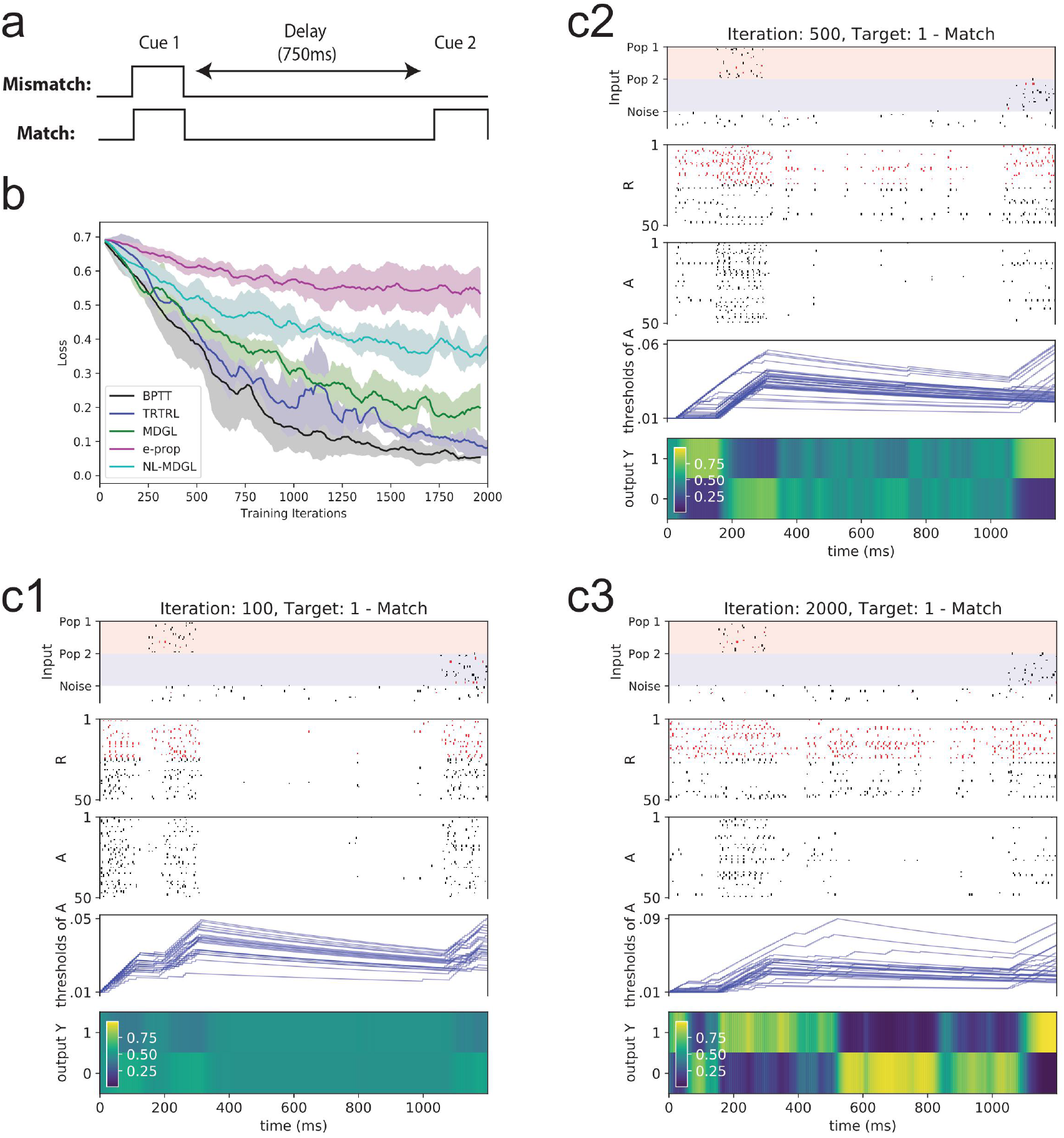
Application of the cell-type-specific modulatory signals to the delayed match-to-sample task. a) Setup of a special case of the delayed match to sample task, where two cue alternatives are represented by the presence and absence of input spikes. b) Learning curves of aforementioned training methods in Figure 2a. Loss is obtained from test data using different realizations of random Poisson input than in training data. Solid lines show the mean and shaded regions show the standard deviation across five runs, with a different weight initialization for each run. Comparing the performance of e-prop with the MDGL method suggests that the addition of cell-type-specific modulatory signals improves learning. c) Network dynamics of an example trial after 100, 500 and 2000 iterations of training using the MDGL method are illustrated in c1, c2 and c3, respectively. To emphasize the change in dynamics over training iterations, we used the same cue pattern for the illustrations. Again, E cells and I cells are color coded using black and red, respectively. For this task, both recurrent units with adaptive threshold (labeled as A) and without (labeled as R) are involved [48]. Threshold dynamics of sample neurons are illustrated. The network makes the correct prediction with greater confidence as training progresses. Figure S5 shows similar results with the nonzero firing representation of the second cue alternative as well.

Figure 5b displays the learning curve for novel (test) inputs for the five plasticity rules described above. We observe that the same general conclusions as for the pattern generation task hold here: TRTRL (blue) and MDGL (green) outperform e-prop (magenta). The performance degrades when we remove the neighborhood specificity of the modulatory signal (NL-MDGL, cyan). Moreover, alignment angles between MDGL, TRTRL, and e-prop are also similar to those of the pattern generation task (Supplementary Tables S1 and S2). We illustrate the network dynamics after training using the MDGL method for 100, 500 and 2000 iterations in Figure 5c1–c3. We observe that as training progresses, the network output decides on the correct prediction with greater confidence, i.e. the output neuron corresponding to the correct target approaches a value of 1. Figure S5 shows that these observations also hold for nonzero firing representation of the second cue alternative.

### Multidigraph learning in RSNNs improves efficiency in an evidence accumulation task

Finally, we study an evidence accumulation task [70, 69], which involves integration of several cues in order to produce the desired output at a later time: an agent moves along a straight path while encountering a series of sensory cues presented either on the right or left side of a track (Figure 6a). Each cue is represented by 40 Hz Poisson spiking input for 100ms and cues are separated by 50ms. After a delay of 850ms when the agent reaches a T-junction, it has to decide if more cues were received on the left or right. Thus, this task requires not only recalling past cues, but also being able to count the cues separately for each side and then process these cues for a reward delivered at a much later time. While e-prop can be used to train a RSNN to solve this task [48], it requires (i) significantly more training iterations than BPTT, and (ii) rapid and stochastic creation/pruning of synapses after each iteration [62]. Therefore, we test our learning rule in Figure 6 to see if the addition of diffuse modulatory signals can indeed bring the learning curve closer to BPTT, without relying on stochastic rewiring.

**Figure 6:**
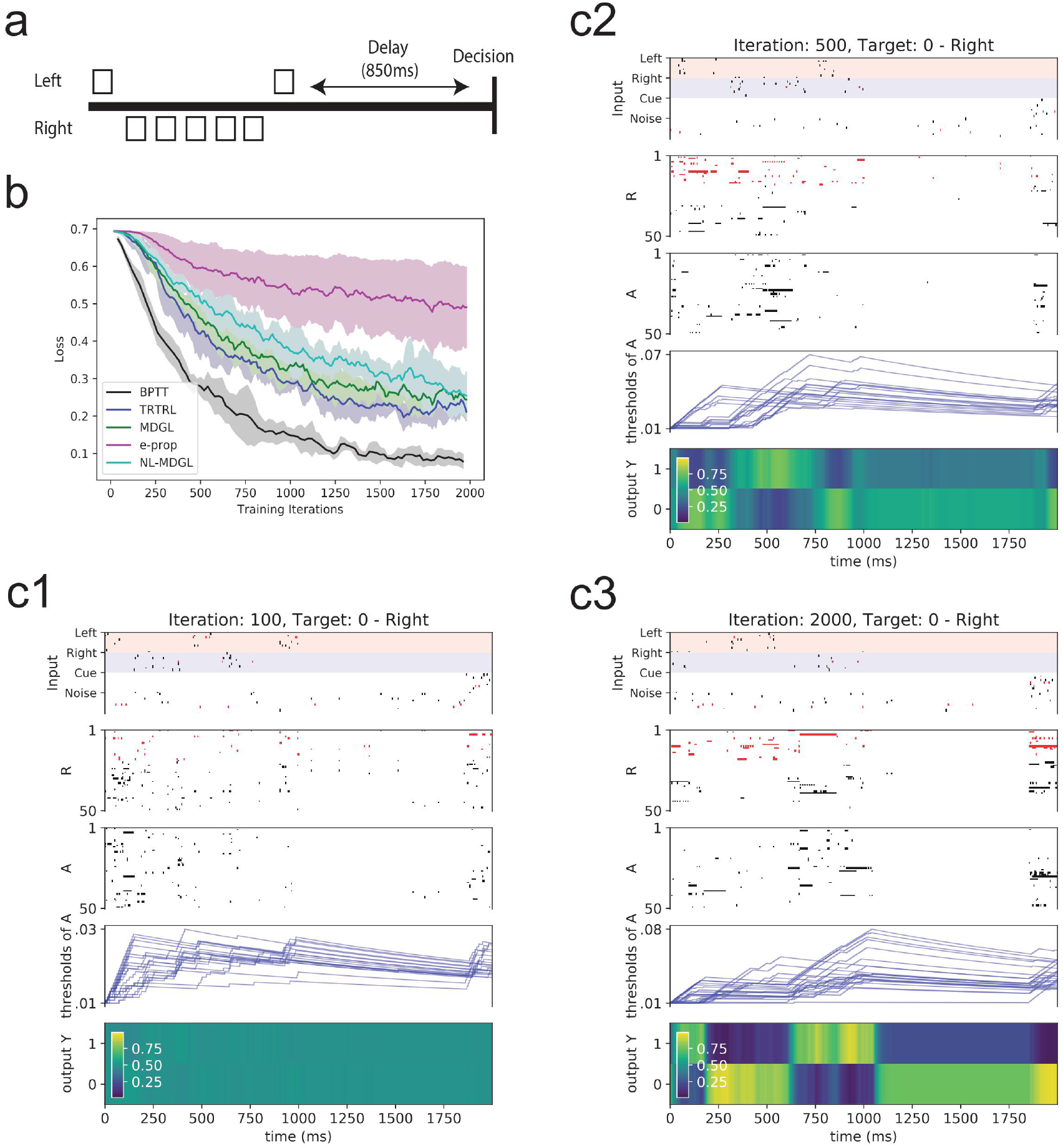
Application to the evidence accumulation task. a) Setup of the task inspired by [69, 70]. b) Learning curves of aforementioned training methods in Figure 2a. Loss is obtained from test data using different realizations of random Poisson input than in training data. Solid lines show the mean and shaded regions show the standard deviation across five runs, with a different weight initialization for each run. Comparing the performance of e-prop with the MDGL method suggests that the addition of cell-type-specific modulatory signals improves learning. c) Network dynamics (Input spikes, recurrent unit spikes and readout) of an example trial after 100, 500 and 2000 iterations of training using the MDGL method are illustrated in c1, c2 and c3, respectively. To emphasize the change in dynamics over training iterations, we used the same target direction for the illustrations. Similar to the previous task, both recurrent units with adaptive threshold (labeled as A) and without (labeled as R) are involved [48], and threshold dynamics of sample neurons are illustrated. The network makes the correct prediction with greater confidence as training progresses. For all methods, results were obtained without using stochastic rewiring, which allows for random formation of new synapses in each experience (Deep R) [62, 63].

Figure 6b and Supplementary Tables S1 and S2 demonstrate that all of our conclusions in the previous two experiments continue to hold in this task: MDGL is closest to BPTT, and NL-MDGL gives relatively degraded performance yet still outperforms e-prop. Figure S10 further shows that gradients approximated by TRTRL and MDGL are more similar to the exact gradients compared to e-prop for the previous two tasks as well as this one. We also observe that the timing of recurrent unit spiking patterns tightly follows that of input cue presentation, suggesting that the network is taking immediate action (“counting”) for each cue. We illustrate the network dynamics after training using the MDGL method for 100, 500 and 2000 iterations in Figure 6c1–c3. We observe that as training progresses, the network output decides on the correct prediction with greater confidence.

While the adaptive firing threshold already provides a form of memory, Figure S6a shows that both threshold adaptation and recurrence are needed for this memory task. (i.e., adaptive threshold alone is not sufficient.) Moreover, Figure S6b shows that this conclusion is not sensitive to the precise fraction of the adaptive population.

### Multidigraph learning in RSNNs produce fast synaptic signaling and slow modulatory signaling

Owing to their diffusive nature and the fact that GPCRs act on significantly longer time scales than receptors for many synaptic neurotransmitters [8], modulatory signaling channels are limited to be slower and smoother (i.e., have lower bandwidth) than direct synaptic channels. Since our model does not explicitly limit the communication bandwidths of either of these channels, comparing the frequency content of these two channels offers an important check for biological plausibility. It also provides a test of our assumption that the summation over *j* in Eq. 3 acts as a smoothing operation, enabling subsequent approximation with diffuse modulatory signaling. Figure 7 demonstrates that the modulatory input indeed has significantly lower frequency content than the synaptic input, for all three tasks studied above.

**Figure 7:**
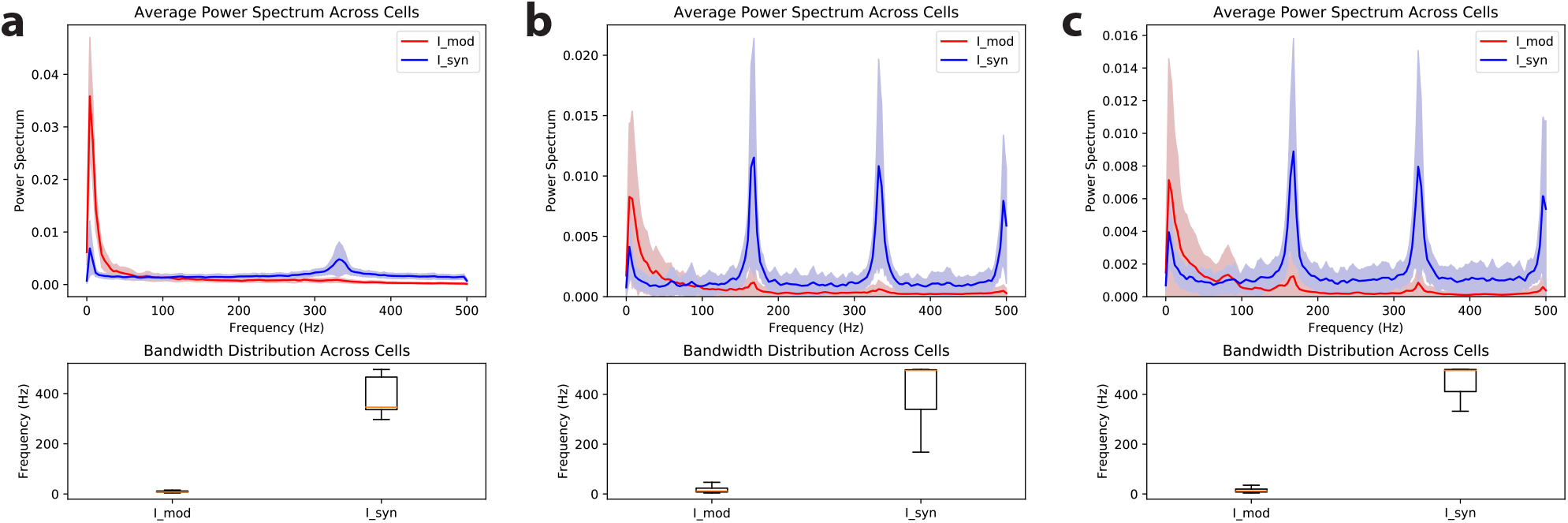
Frequency distribution of modulatory versus synaptic inputs. Comparing the power spectrum and bandwidth distributions between modulatory and synaptic inputs for a) pattern generation, b) delayed match to sample and c) evidence accumulation tasks. In the top panel, the solid lines denote the average and shaded regions show the standard deviation of power spectrum across recurrent cells. In the bottom panel, box plots show the minimum, lower quartile, median, upper quartile and maximum bandwidth across the cells. Here, the bandwidth is quantified by the 3dB frequency, where the power halves compared to the peak. Nyquist’s theorem dictates that the simulation interval of 1ms limits the maximum frequency to 500Hz. Modulatory input is the total cell-type-specific modulatory signals detected by each cell *p*, defined as 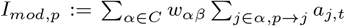 (Eq. 5). Synaptic input is the total input received through synaptic connections by each cell *j*, defined as 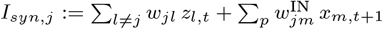 (Eq. M1). The naive implementation of the modulatory signal, *a*_*j,t*_, depends on future errors unless the output is not leaky (Eq. M12 in Methods, Supplementary Note 1). Thus, we repeated the spectral analysis for the online implementation of modulatory signaling in Eq. S3 and observed similar conclusions (Figure S4). Here, the plots are obtained at the end of training (after 500 iterations for pattern generation and 2000 iterations for delayed match to sample and evidence accumulation tasks), but similar trends are observed at other training snapshots.

In sum, our results on the slow timescales of cell-type averaged signaling – together with the gradient alignment results mentioned above (presented in supplementary materials) – help to illustrate why modulatory signaling that is non-specific across both space and time can nonetheless lend significant improvement to network learning curves. This form of signaling removes the need for specifically tuned, reciprocal physical contacts and signaling among cell pairs and hence biologically implausible features of anatomical organization that plague solutions to the credit assignment problem via pairwise synaptic communication alone.

## Discussion

In this paper, we presented a normative theory of temporal credit assignment and an associated biologically plausible learning rule where neurons are allowed to communicate via not only the synaptic connections but also a secondary, non-specific “modulatory” channel. In particular, we explained how the recent observation of wide-spread and cell-type-specific modulatory signaling [8], when integrated with synaptic networks to interconnect cortical neurons, can promote efficient learning. We demonstrated that the associated biologically plausible learning rule achieves performance close to that of ideal, but biologically unrealistic, rules on multiple *in silico* learning tasks – pattern generation, delayed match-to-sample, and evidence accumulation – that require temporal credit assignment over timescales spanning seconds. These experiments used sparsely and recurrently connected neural networks of sign-constrained spiking neurons, capturing some of the well-studied aspects of brain architecture and computation [60, 61]. All simulation results were obtained without using rapid stochastic rewiring, which allows for random formation of new synapses in each experience [62, 63]. Thus, learning with local modulatory signals can relax the need for rapid and random creation/pruning of synapses. While our demonstrations use the supervised learning paradigm, the same machinery can be easily applied to reinforcement learning tasks [48].

The existence of multiple directed connections between neurons suggests a *multidigraph* view of brain connectivity, in which each neuron is connected by multiple different connection types (Figures 1 and 2b) [8]. Therefore, we call our learning paradigm, multidigraph learning. MDGL begins with the same basic structure as the RFLO [47] and e-prop [48] learning rules, but adds the cell-type-specific local modulatory network to achieve a better approximation to ideal synaptic weight updates. While we did not constrain the temporal dynamics of this secondary signaling mechanism, we found that the cell-type-specific broadcast channel communicated at significantly lower frequencies in all our experiments, in agreement with well-known properties of diffusive modulatory signaling.

Our model posits that top-down learning signals can influence the secretion of cell-type-based modulatory signals (Eq. 7 and Supplementary Note 2). As one possible mechanism for this, dopamine can alter neuronal firing [71], which can then impact neuropeptide release in an activity-dependent manner [39]. This finding predicts that cell-type-specific signaling carries information about top-down learning signals, so their levels can reflect the learning progress. Indeed, our computational experiments suggest that the level of modulatory input decreases over training iterations and sharply rises in response to changes in task condition (Figure S7). Rapidly maturing technologies on *in-vivo* monitoring of neuromodulators [72, 73, 41] suggest that such predictions of our model can be tested experimentally.

The nature of “intermediate” cells [9], whose phenotypes appear to be a mixture of “pure” cell types is a key problem in cell types research. Our findings explain the existence of such phenotypes from a connectivity perspective: while average connectivities between types *w*_*αβ*_ remain relatively constant during training, connectivities of individual cells can deviate significantly from those averages (Figure S8). We also hypothesized a link between abstract cell type-based connectivities and modulatory receptor efficacies. From this perspective, how tightly the individual synaptic weights and cell-type-specific (but not neuron specific) receptor efficacies should be coupled (i.e., co-adaptation) may be explored in future work. Figure S9 suggests that the effect of imprecise GPCR efficacies (*w*_*αβ*_) on the performance is task-dependent, and a wide range of fixed *w*_*αβ*_ values can enable effective learning in some, but not all, tasks.

As a proof of concept, we tested a minimal implementation of MDGL: there are just two local modulatory types mapping to the two main cortical cell classes (E and I), and the fine-grained E cell subtypes (those with and without firing threshold adaptation) were grouped into one modulatory type. Even the minimal implementation led to a marked improvement in learning efficiency compare to existing biological learning rules. However, brain cells are extremely diverse [74,75,76,77,78,79] with a matching diversity in the expression of peptidergic genes; at least 18 NP precursor and 29 NP-GPCR genes were reported to have widespread and cell-type-specific expression in the mouse cortex [7]. Therefore, further studies can investigate how MDGL scales to harder tasks and examine the interplay of task complexity and cell diversity. A related point is that, here, we did not study the potential diversity of time scales across different types of NP signals [39]. Future work may investigate the role of distinct timescales in local modulatory signaling.

Our MDGL learning rule assumes that cell-type-based modulatory signals diffuse locally such that they are registered only by local synaptic partners of the source neuron. On the other hand, NL-MDGL assumes that these signals diffuse to all cells of a given type in the network. The biological reality may lie somewhere in between. Studying the extent of the local diffusion hypothesis, where neuropeptide signals act on both synaptic partners and nonsynaptic partners in the vicinity [39] may suggest new learning rules in the future and deepen our understanding of their underlying mechanisms.

Learning rules often explicitly optimize a loss function, and the direction of the steepest descent in the loss function can be identified from the exact gradient, when available. Rules that follow this exact gradient, RTRL and BPTT, are well established, but are not biologically plausible and have unmanageably vast memory storage demands. However, a growing body of studies have demonstrated that learning rules that only partially follow the gradient, while alleviating some of these problems of the exact rules, can still lead to desirable outcomes [80, 81]. An example is the seminal concept of feedback alignment [64], which rivals backpropagation on a variety of tasks even using random feedback weights for credit assignment. In addition, approximations to RTRL have been proposed [82, 83, 84, 85] for efficient online learning in RNNs. Reference [47] also derived a *O*(*N* ^2^) rule that maintained only a rank-2 tensor for training rate-based RNNs by fully truncating the inter-cellular dependencies involving cells other than pre-and postsynaptic neurons. Instead of full truncation, SnAp-*n* [54, 55] *stores the Jacobian* 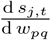 for all *j* influenced by *w*_*pq*_ within *n* time steps. SnAp-1 is effectively the truncation in [47] and the performance of SnAp-*n* increases with *n*, resonating with our improved performance when more terms of the exact gradient are included. However, SnAp-*n* needs to maintain a triple tensor for *n* ≥ 2. While our approximation is similar to SnAp-2, it only needs to store the double tensor 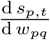 since we further constrain inter-cellular signaling via diffuse pathways organized by cell types. Thus, our model further advances the efficacy of approximated gradient-based learning methods and continues the line of research in energy-efficient on-chip learning through spike-based communications [86, 87]. Such efficient approximations of the gradient computation can be especially important as artificial networks become ever larger and are used to tackle ever more complex tasks under both time and energy efficiency constraints.

Our work fits under the wide umbrella of neoHebbian multi-factor learning [21, 26]. This theory posits additional modulatory factors, which combine with factors based on pre- and postsynaptic activity to determine synaptic plasticity. Molecules that could keep a memory trace of classical Hebbian activity include calcium ions and activated CaMKII enzymes [88] although the biological underpinning remains enigmatic. The additional modulatory factors could arise from a diverse set of signals found in the brain. Most existing theoretical models conceive these factors as top-down learning signals coming from distant brain regions, such as dopamine, noradrenaline and neural firing [89]. Here, we took top-down learning signals to be cell-specific rather than global, which is justified in part by recent reports that dopamine signals [69] and error-related neural firing [89] can be specific to a population of neurons [48]. On the other hand, this specificity could be removed by random feedback alignment [64] or via an approach similar to our cell-type-specific weight approximation in Eq. 4 (Supplementary Note 2). In addition to top-down learning signals, we add further modulatory factors, released from within the recurrent network performing the computation at hand. These signals represent locally secreted modulatory signals, such as those carried by neuropeptides, released from nearby cells of a given type. Such signalling is abundantly present in the brain but had yet to be interpreted in terms of the credit assignment problem. Similar to dopaminergic regulation of plasticity [71], neuropeptides have also been shown to modulate plateau potential, which could alter the amount of plasticity [90].

Our work suggests that multiple cell-type-specific, diffuse and relatively slow modulatory signals should be considered as possible bases for credit assignment computations. Though inspiration for the present work came primarily from new transcriptomic data on local NP signaling in neocortex [7, 8], it is quite possible that other cell-type-specific neuromodulators could likewise contribute to credit assignment. Many of these alternative agents act, as do NPs, via GPCRs (e.g., the monoamines, amino acids, acetylcholine and endocannabinoids, as noted above), but our multidigraph template might even apply to other neuronally secreted neuromodulators, such as the neurotrophins and cytokines, that act via different classes of receptor [91, 92]. While experimental tests of such hypotheses have not seemed feasible up until now, emerging methods for genetically addressed measurement of various neuromodulatory signals in specific cell types [7, 93, 94] are now bringing the necessary critical tests within reach (e.g. [41]).

## Methods

### Neuron Model

We consider a discrete-time implementation of RSNNs. The network, as shown in Figure 3a, denotes the observable states, i.e. spikes, as *z*_*t*_ at time *t*, and the corresponding hidden states as *s*_*t*_. For LIF cells, the state *s*_*t*_ corresponds to membrane potential and the dynamics of those states are governed by

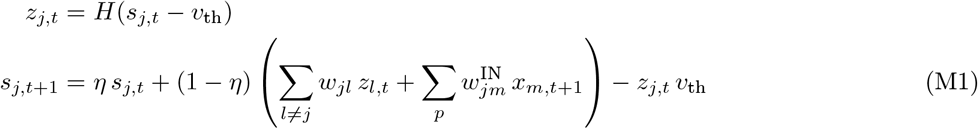

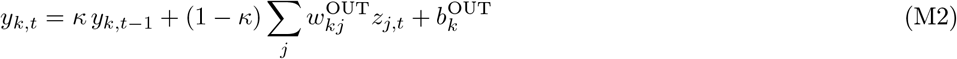

where *s*_*j,t*_ denotes the membrane potential for neuron *j* at time *t, v*_th_ denotes the spiking threshold potential, 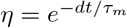 denotes the leak factor for simulation time step *dt* and membrane time constant *τ*_*m*_, *w*_*lj*_ denotes the weight of the synaptic connection from neuron *j* to *l*, 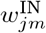 denotes the efficacy of the connection between the input neuron *m* and neuron *j*, and *H* denotes the Heaviside step function. For the output, 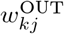 denotes the efficacy of the connection from neuron *j* to output neuron *k*, and 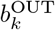 denotes the bias of the *k*-th output neuron.

Following references [63, 48], which implemented adaptive threshold LIF (ALIF) units [95] and observed that this neuron model improves computing capabilities of RSNNs relative to networks with LIF neurons, we also include ALIF cells in our model. In addition to the membrane potential, ALIF cells have a second hidden variable, *b*_*t*_, governing the adaptive threshold. The spiking dynamics of both LIF and ALIF cells can be characterized by the following set of equations:

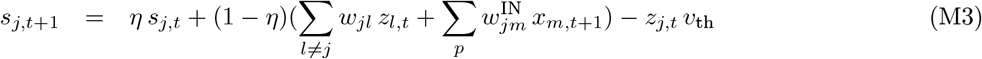

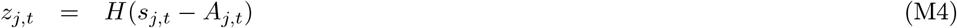

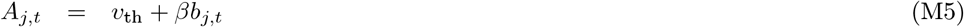

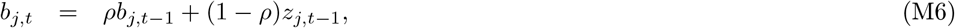

where the voltage dynamics in Eq. M3 is the same as Eq. M1. A spike is generated when the voltage *s*_*j,t*_ exceeds the dynamic threshold *A*_*j,t*_. Parameter *β* controls how much adaptation affects the threshold and state *b*_*j,t*_ denotes the variable component of the dynamic threshold. The decay factor *ρ* is given by 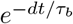 for simulation time step *dt* and adaptation time constant *τ*_*b*_, which is typically chosen on the behavioral task time scale. For regular LIF neurons without adaptive threshold, one can simply set *β* = 0.

### Network output and loss function

Dynamics of leaky, graded readout neurons was implemented as 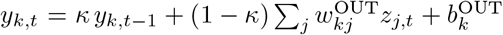. The (1 − *κ*) factor in the second term was dropped in writing for readability but kept during the actual implementation. Here, *κ ∈* (0, 1) defines the leak and 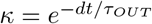 for output membrane time constant *τ*_*OUT*_. We provide the online implementation for this readout convention in Supplementary Note 1.

We quantify how well the network output matches the desired target using error function *E*. For regression tasks such as pattern generation, we use 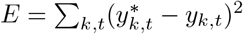 given time-dependent target 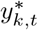 For classification tasks such as delayed match to sample and evidence accumulation, 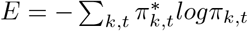 with one-hot encoded target 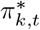 and predicted category probability *π*_*k,t*_ = softmax_*k*_(*y*_1,*t*_, …, *yN*_*OUT*_,*t*) = exp(*y*_*k,t*_)*/* Σ_*k*_*’* exp(*y*_*k*_*’*,*t*). We provide all simulation and training parameters in Supplementary Note 3.

### Notation for Derivatives

There are two types of computational dependencies in RSNNs: direct and indirect dependencies. For example, variable *w*_*pq*_ can impact state *s*_*p,t*_ directly through Eq. M1 as well as indirectly via its influence through other cells in the network. We distinguish direct dependencies versus all dependencies (including indirect ones) using partial derivatives (*∂*) versus total derivatives (*d*).

### Differentiation in RSNNs

As mentioned, we study iterative adjustment of all synaptic weights using gradient descent on loss *E*:

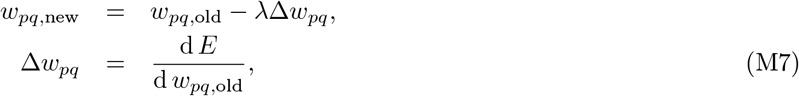

where *λ* denotes the learning rate, and the gradient of the error with respect to the synaptic weights must be calculated. As explained before in Results, gradient-based learning rules, BPTT and RTRL, calculate this by unwrapping the RSNN dynamics over time. While the two algorithms yield equivalent results, we focus our analysis on RTRL, since its computational dependency graph of states is causal. Using RTRL, we factor the error gradient as follows:

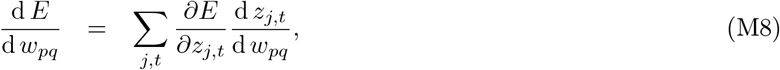

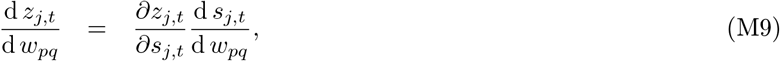

following the derivation notation explained above. The factor 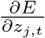 in Eq. M8 is related to the top-down learning signal 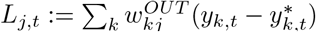 [48]. Supplementary Notes 1, 2 show that the leak term of the output neurons makes these two terms different, and derives an online implementation that uses *L*_*j,t*_. For readability, however, we simply refer to 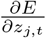 as top-down learning signal in the main text, leaving the detailed expansion of derivatives to the supplementary materials.

We now discuss the second factor in Eq. M8, i.e. 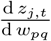. This is expanded in two factors in Eq. M9. The first factor, 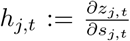 is problematic to compute for spiking neurons due to the discontinuous step function *H* in Eq. M3, whose derivative is not defined at 0 and is 0 everywhere else. We overcome this issue by approximating the decay of the derivative using a piece-wise linear function [63, 86, 96, 87]. Here, the pseudoderivative *h*_*j,t*_ is defined as follows:

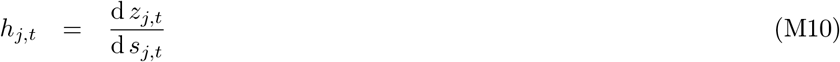

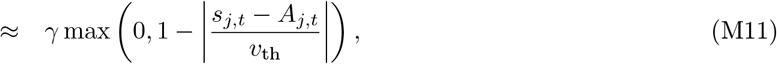

The dampening factor *γ* (typically set to 0.3) dampens the increase of backpropagated errors in order to improve the stability of training very deep (unrolled) RSNNs [63]. Throughout this study, refractoriness is implemented as in [48], where *h*_*j,t*_ and *z*_*j,t*_ are fixed at 0 after each spike of neuron *j* for 2 to 5ms.

The last factor, 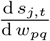, accounts for both spatial and temporal dependencies in RSNNs and can be obtained recursively using Eq. 1. As seen from Eq. 1 and explained in Results, this factor poses key issues to biological plausibility and computational cost.

### Cell-type-specific signaling implementation

As introduced in Eq. 5, activity-dependent modulatory signal emitted by neuron *j* at time *t*, an important component of MDGL, is defined as

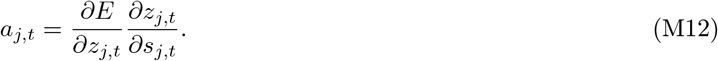

As defined, *a*_*j,t*_, is a package of two components: 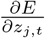, which is referred to as the top-down signal [48], and *h*_*j,t*_ = *∂z*_*j,t*_*/∂s*_*j,t*_, which is the pseudo-derivative of spiking activity as a function of cell *j*’s membrane potential explained above.

### Eligibility Trace Implementation

As introduced in Eq. 5, eligibility trace, another important component of MDGL, is defined as

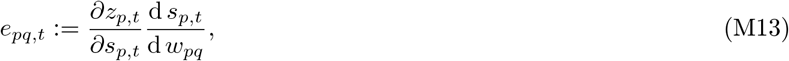

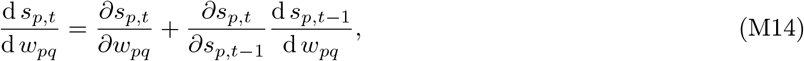

where Eq. M14 follows directly from Eq. 2. 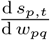 can be obtained recursively and is referred to as the eligibility vector [48]. *e*_*pq,t*_ keeps a fading memory of activity pertaining to presynaptic cell *q* and postsynaptic cell *p*.Here, we briefly explain its implementation by expanding the factors in Eqs. M13 and Eq. M14 for both LIF and ALIF cells.

For LIF cells, there is no adaptive threshold so the hidden state consists only of the membrane potential. Thus, we have factors 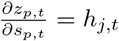 with pseudo-derivative *h*_*j,t*_ defined in Eq. M10, 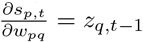 and 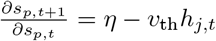 following Eq. M1.

For ALIF cells, there are two hidden variables so the eligibility vector is now a two dimensional vector 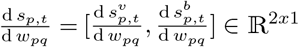 pertaining to membrane potential *v*_*p,t*_ and adaptive threshold state *b*_*p,t*_. Following Eq. M3, one can obtain factors 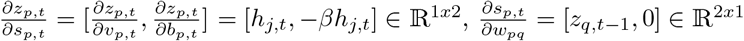 and 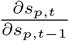 is now a 2-by-2 matrix:

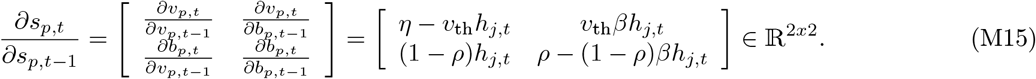

Thus, the eligibility trace *e*_*pq,t*_ would be scalar valued regardless of the dimension of the eligibility vector.

### Firing Rate Regularization

In addition to accuracy optimization described above, we added a firing rate regularization term *E*_*reg*_ to the loss function to ensure sparse firing [48]:

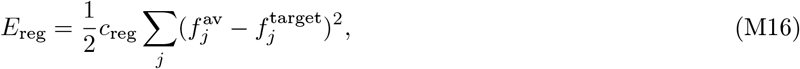

where 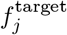 and 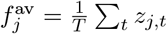 are the desired and actual average firing rate for cell *j*, respectively, and *c*_reg_ is a positive coefficient that controls the strength of the regularization.

## Acknowledgements

We are grateful to Guillaume Lajoie, James Murray, and Scott Owen for helpful feedback on the manuscript. We also wish to thank the Allen Institute for Brain Science founder, Paul G Allen, for his vision, encouragement and support. Helena Liu is supported by Natural the Science and Engineering Research Council (NSERC) Postgraduate Scholarships -Doctoral (NSERC PGS-D) program. This work was facilitated through the use of advanced computational, storage, and networking infrastructure provided by the Hyak supercomputer system at the University of Washington.

## Supplementary Materials

## Supplementary Figures

**Figure S1:**
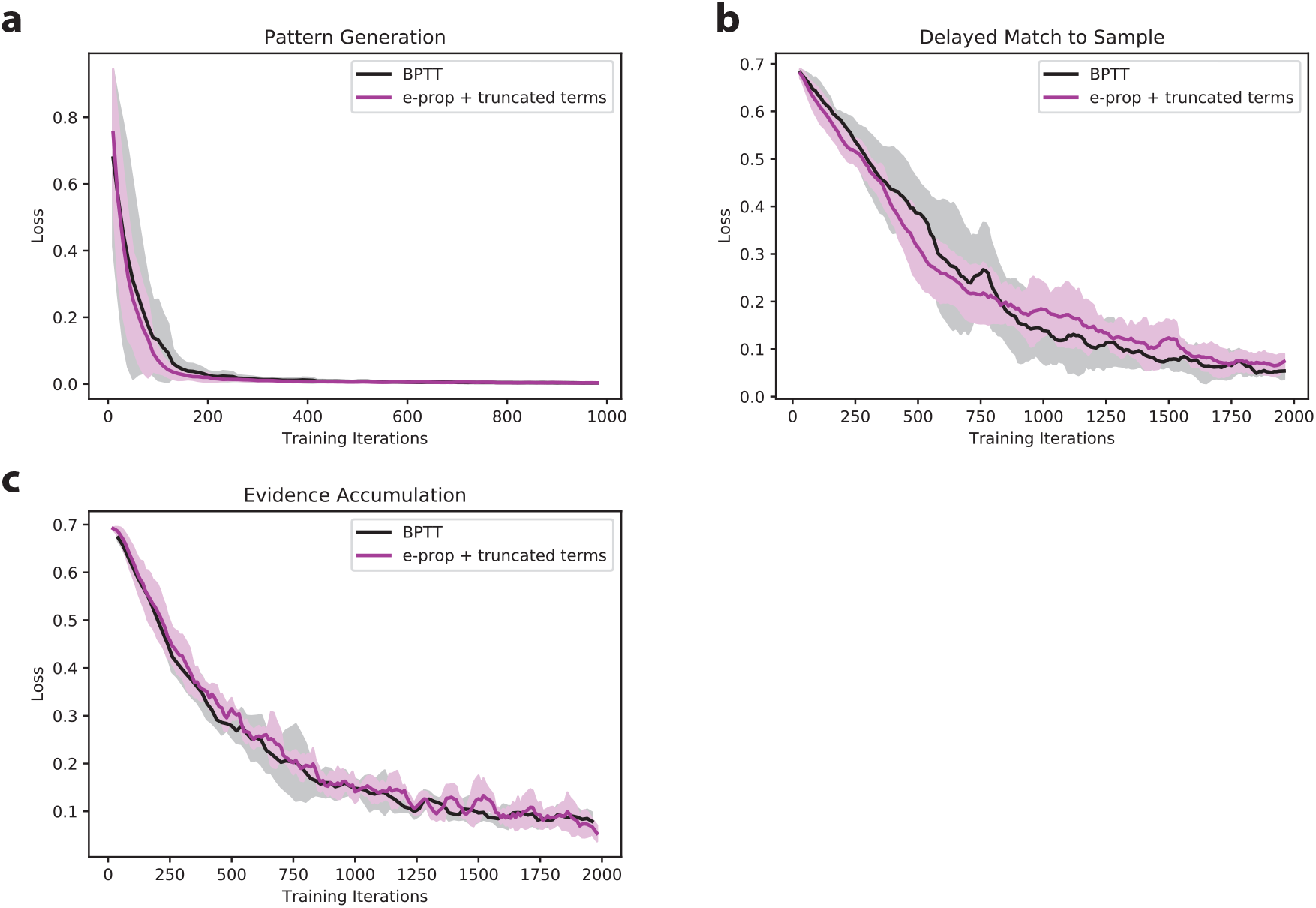
Checking e-prop implementation – recovering ignored terms recovers the performance of BPTT. As a sanity check, learning curves are plotted for e-prop plus all the truncated terms (see Eq. 1)to verify that the resulting learning rule recovers the performance of BPTT. The check is applied to a) pattern generation, b) delayed match to sample and c) evidence accumulation tasks. Solid lines show the mean averaged across five runs and shaded regions show the standard deviation. For all tasks, the learning curves do not differ significantly, suggesting the e-prop implementation is accurate.

**Figure S2:**
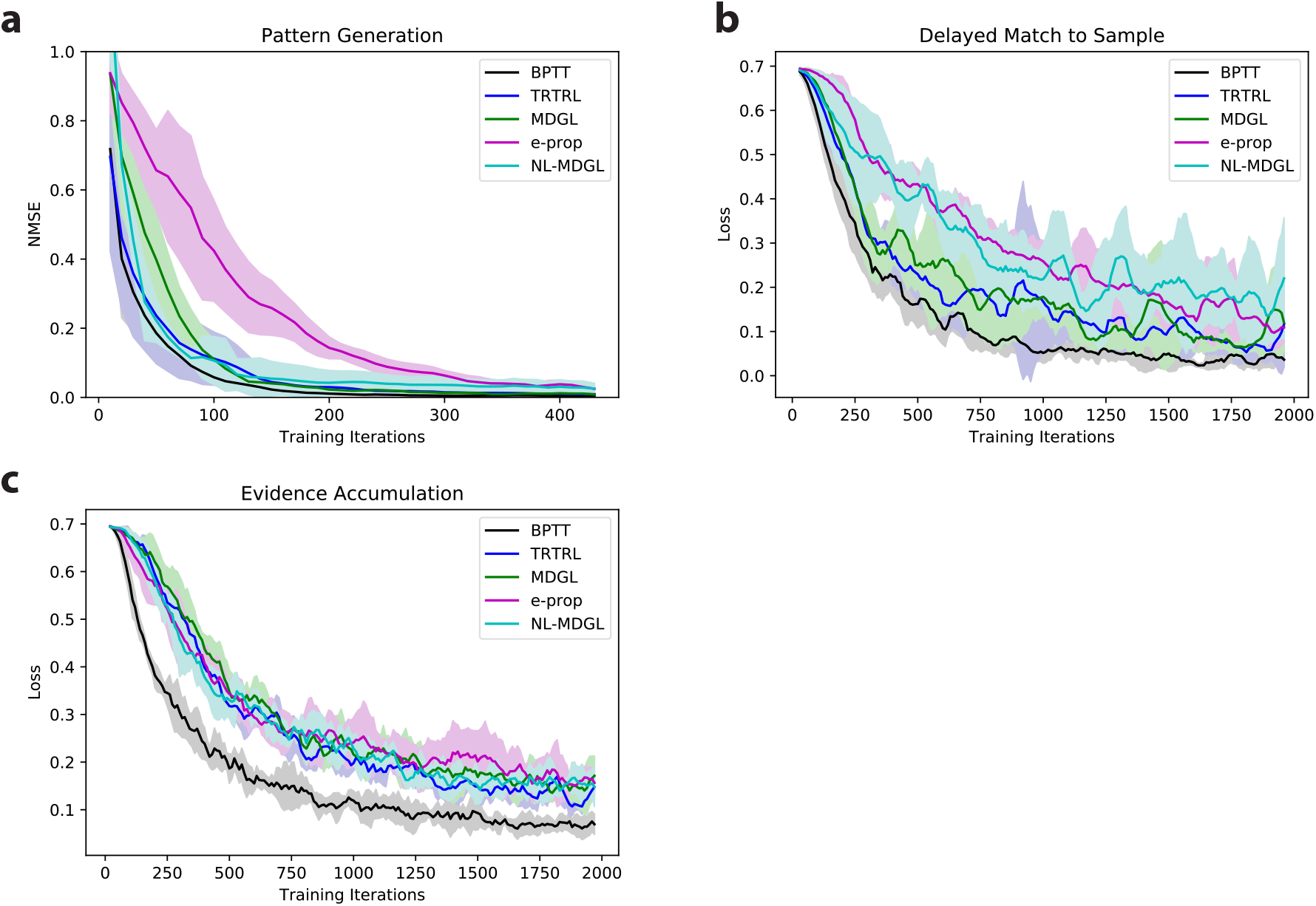
Learning performance of RSNNs with 30% connection probability. Learning curves for training methods in Figure 2a are illustrated for the following tasks: a) pattern generation, b) delayed match to sample and c) evidence accumulation tasks. The learning curve gap here between e-prop and MDGL is narrower than that of the network with 10% connection probability shown in the main text. This suggests that MDGL is more helpful under sparse scenarios. As noted in the main text, connection sparsity is widely observed in the brain [60]and has various computational advantages [97, 98].

**Figure S3:**
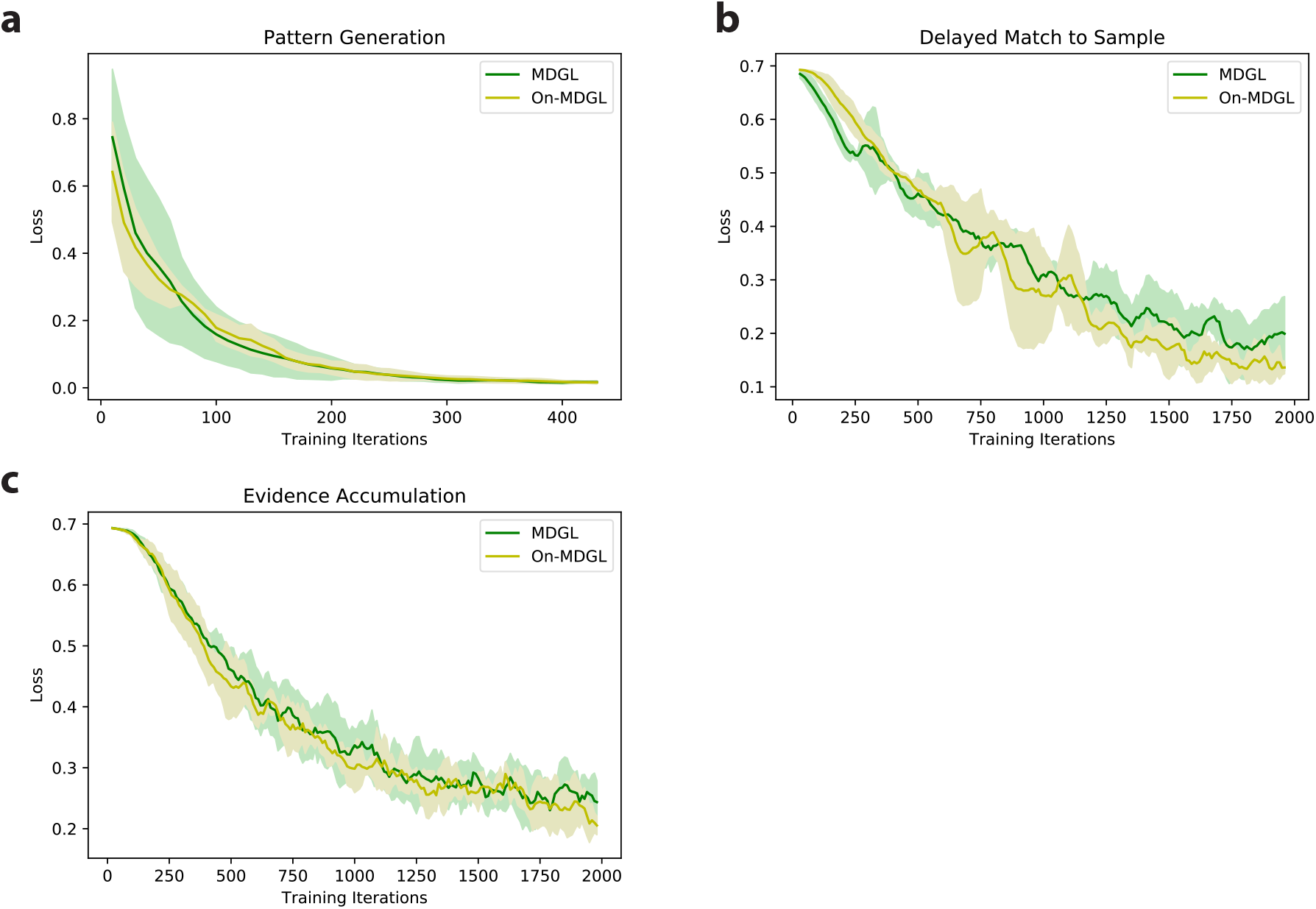
No significant degradation in performance observed for the online approximation of MDGL in Eq. S3. For outputs with a leak term defined in Eq. M2, 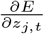 depends on future errors and an approximation is introduced in Eq. S3for online implementation of MDGL. To check if this approximation leads to significant degradation in performance, learning curves are plotted for a) pattern generation, b) delayed match to sample and c) evidence accumulation tasks. For all tasks, there is no significant deviation in learning curves between MDGL and the online approximation (On-MDGL).

**Figure S4:**
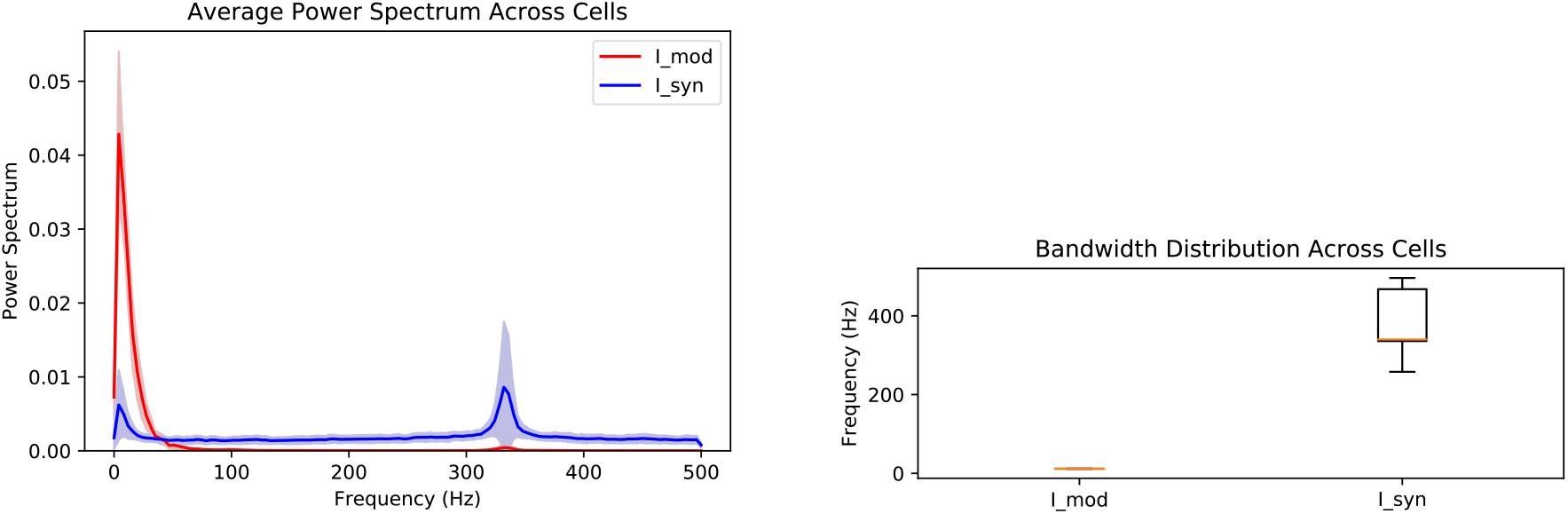
Slowness in modulatory signaling for the online approximation of MDGL. We repeat the spectral analysis in Figure 7, but for the online implementation of local modulatory input in Eq. S3, i.e. 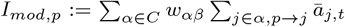 with *ā* _*j,t*_ defined in Eq. S3. The observations here match those of Figure 7, where modulatory input is significantly slower than synaptic input. We note that the analysis here is done on the pattern generation task only, because for the other two tasks, the error signal is not available until the end of the trial, making the modulatory input too short (see Eq. S3)for any meaningful spectral analysis. We expect this “slowness” of modulatory signaling to generalize as the modulatory input is a weighted summation of slow changing leaky outputs and low-pass filtered activity *h*_*j,t*_.

**Figure S5:**
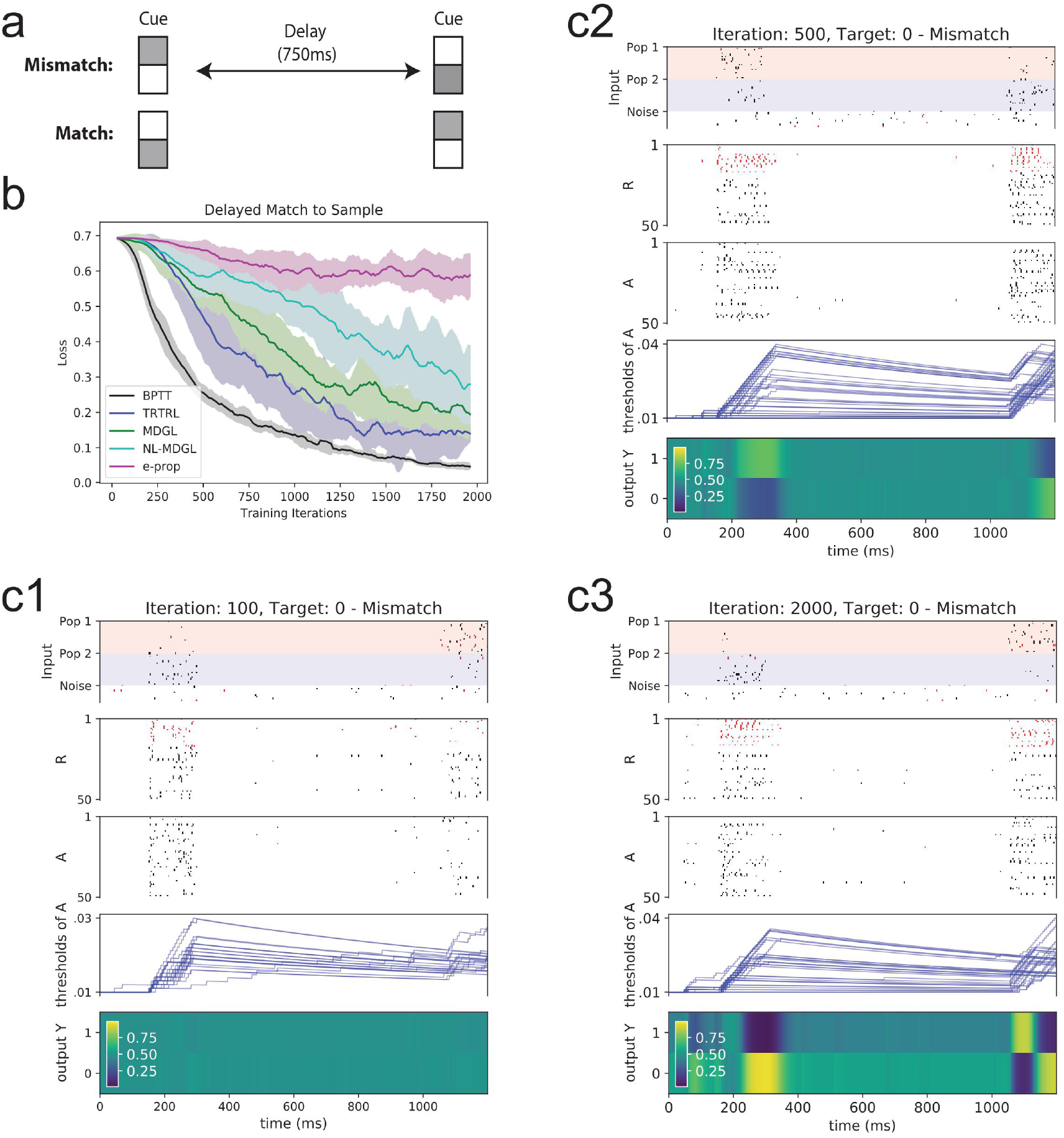
Similar observations for alternative task parameters in the task of Figure 5. The delayed match-to-sample task in Figure 5 is repeated here, but with nonzero firing rates for the second cue alternative. As before, two input populations take on two different firing statistics to represent the two cue alternatives, and the agent is tasked with determining if the cue presented before and after the delay period correspond to the same cue alternative. Here, the rates of these two populations provided in Supplementary Note 3. The plotting conventions here are the same as those of Figure 5, except that a larger network is used (see Supplementary Note 3) so for better readability, only 50 units are selected for the raster plots. The same conclusion as Figure 5 is observed here: comparing the performance of e-prop with the MDGL method suggests that the addition of cell-type-specific modulatory signals expedites the learning curve; the network makes the correct prediction with greater confidence as training progresses.

**Figure S6:**
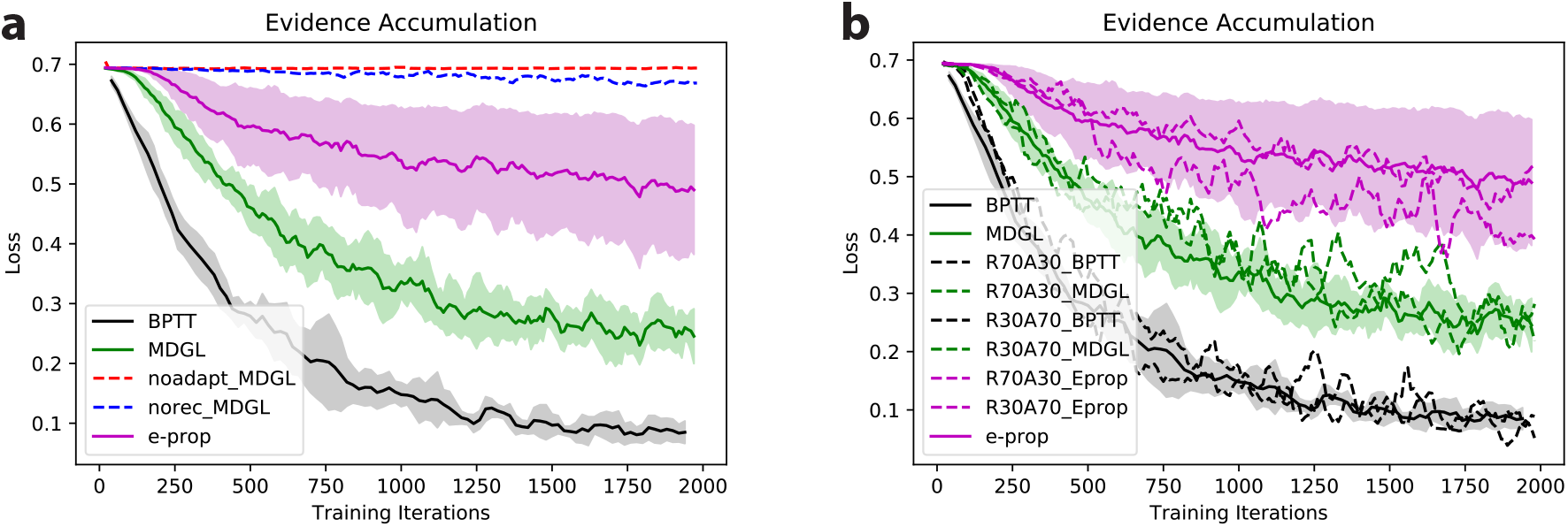
Threshold adaptation analysis for the evidence accumulation task in Figure 6. a) Both threshold adaptation and recurrence are needed for successful completion of the task. MDGL trained on a network without ALIF cells (red) or with recurrent connections removed (blue) shows little decrease in loss function. b) Simulating with 70 LIF to 30 ALIF cells (R70A30) as well as 30 LIF to 70 ALIF cells (R30A79) led to similar ordering of performance for different learning methods as the default (50 LIF to 50 ALIF cells). Here, dashed lines represent individual runs.

**Figure S7:**
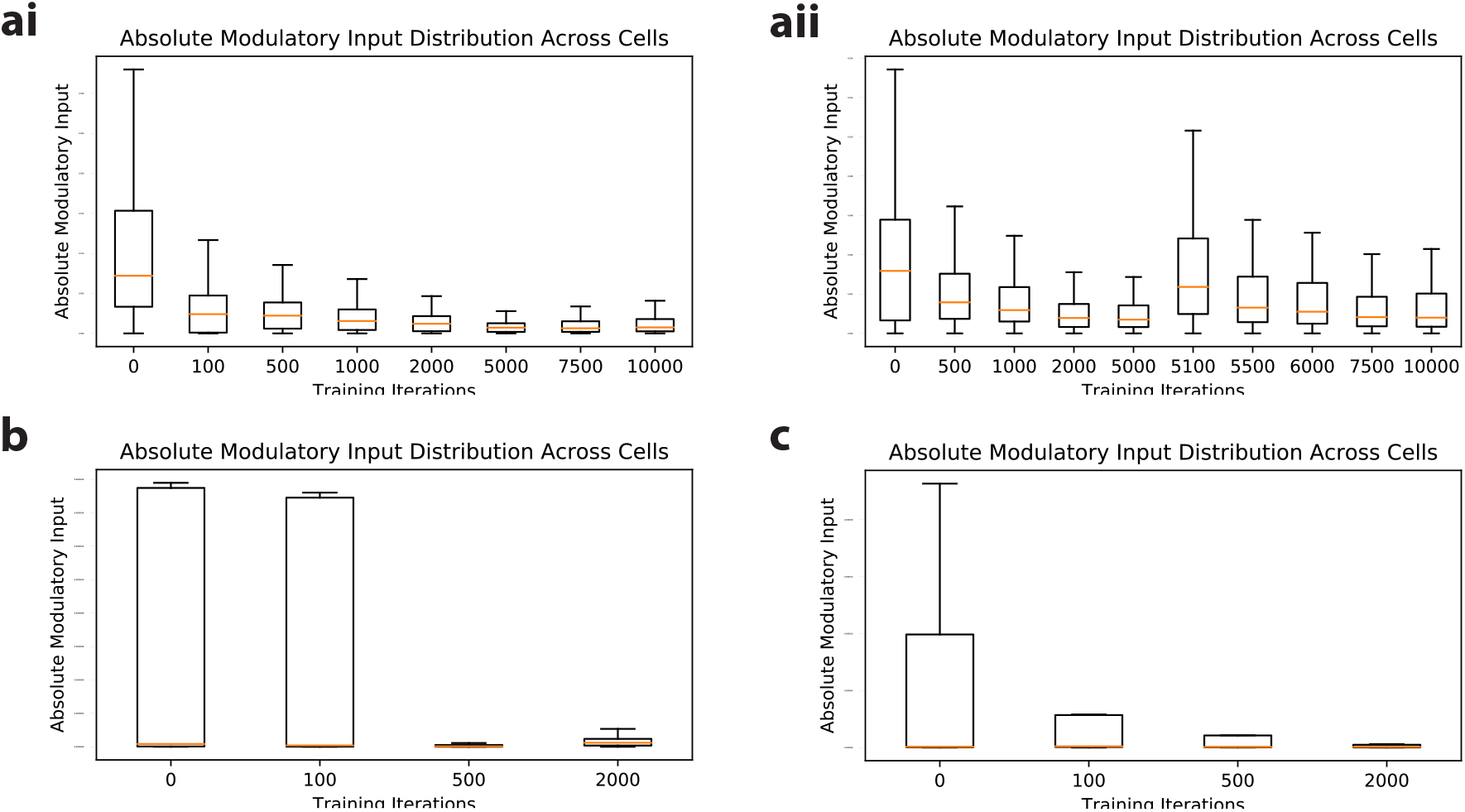
Cell-type-specific modulatory signaling level decreases over training iterations. Box plots for absolute cell-type-specific modulatory input distribution across cells show that modulatory signalling drops over training iterations for ai) pattern generation, b) delayed match-to-sample and c) evidence accumulation tasks. In aii), the target for the pattern generation task was changed after 5000 iterations, which resulted in a rapid increase in modulatory input immediately after the change, and a progressive decrease as training continues. Here, absolute modulatory input for cell *p* is defined as 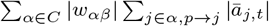 (*ā* _*j,t*_ is defined in Eq. S3),where the absolute value is taken because we are interested in the magnitude of the signaling level.

**Figure S8:**
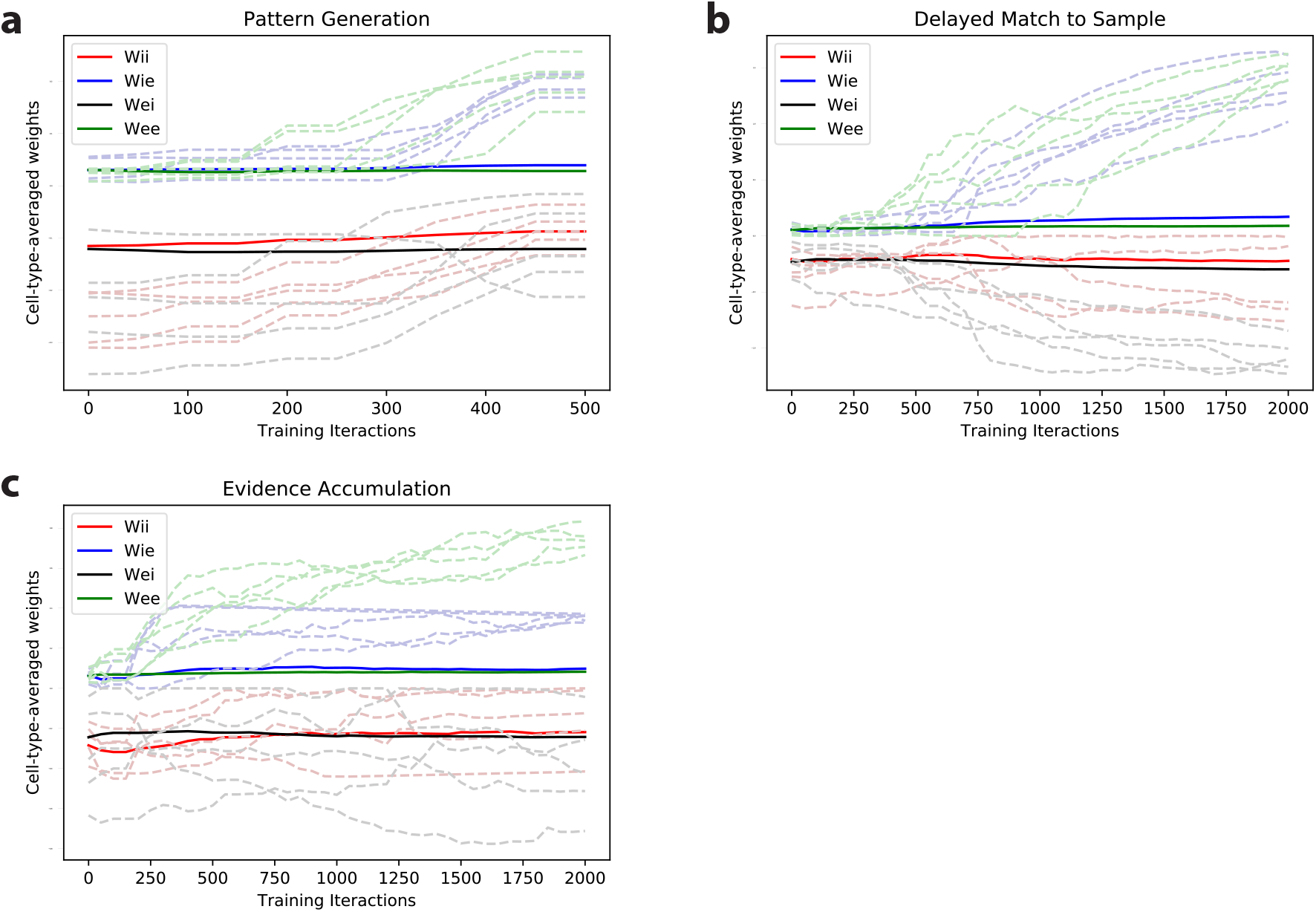
Cell-type-specific signaling gain drifts slowly over training. The four cell-type-specific signaling gains for MDGL with two cell-types, i.e. *w*_*αβ*_ in Eq. 4 and Eq. 5 with *α, β ϵ{E, I}*, are illustrated in solid lines for the three tasks investigated. Several sample individual weights with large changes are illustrated in faint dashed lines. Because we implemented cell-type-specific signaling gain using weight averages, it is not surprising that they drift over training as weights adapt. This drift, however, is slow compared to how fast some individual weights can change.

**Figure S9:**
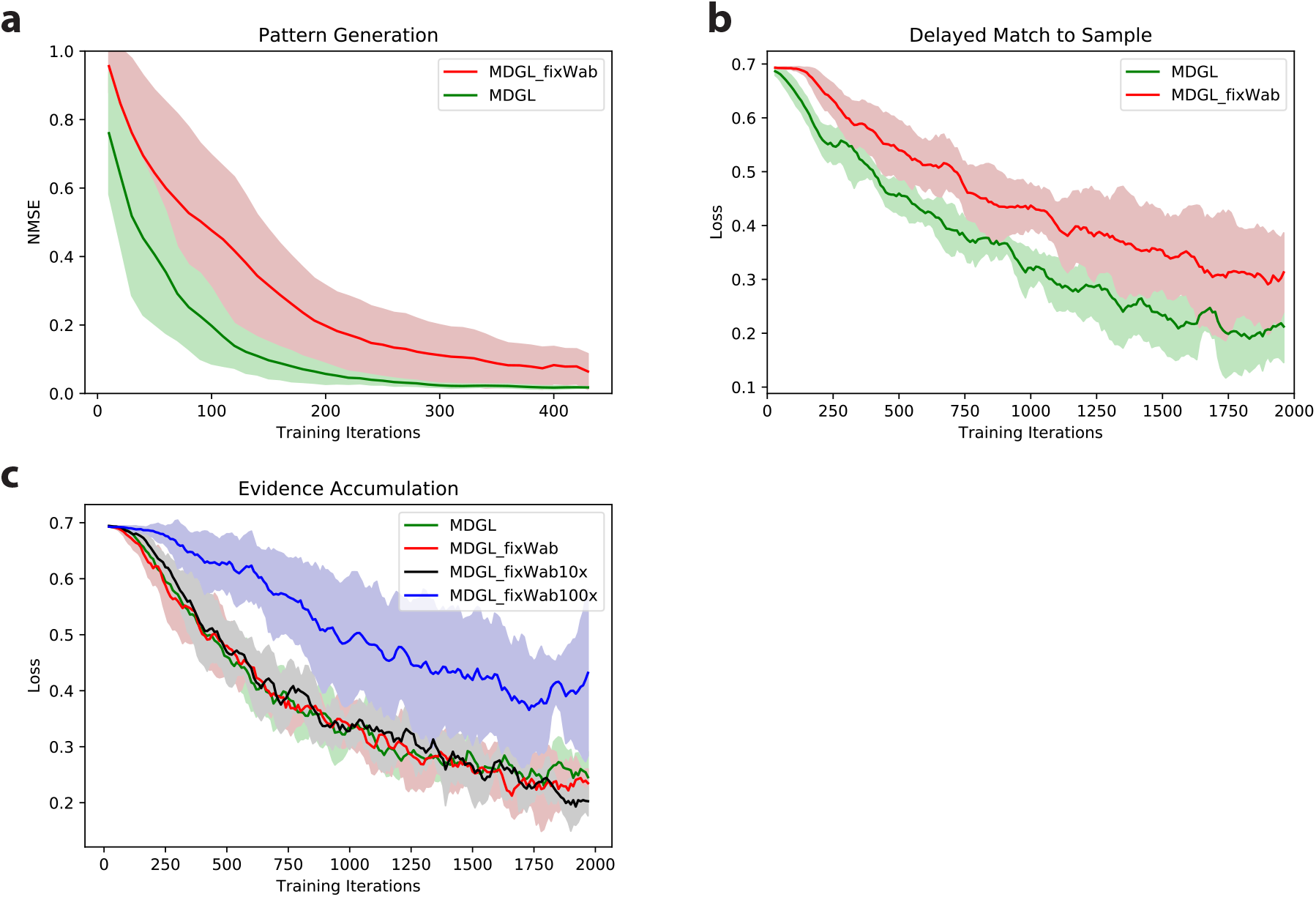
Comparing the learning curves for default MDGL versus MDGL using fixed cell-type-specific gain *w*_*αβ*_. This comparison is done for a) pattern generation, b) delayed match-to-sample and c) evidence accumulation tasks. Here, the magnitude of each *w*_*αβ*_ (for *α ∈ {I, E}, β ∈ {I, E}*) is randomly generated using a Gaussian random variable with zero mean and variance of 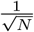 upon each initialization and fixed throughout the training. The sign of *w*_*αβ*_ is taken to be the sign of its presynaptic group. We observe degradation in performance for the pattern generation task and delayed match-to-sample tasks using this fixed random *w*_*αβ*_ (labeled as MDGL_fixWab in each panel), but not for evidence accumulation task. Multiplying this random *w*_*αβ*_ by a factor of 100 (labeled as MDGL_fixWab100x) pushes the network outside of an efficient operating range for the evidence accumulation task, but the training for this task appears to be robust even when the randomly generated *w*_*αβ*_ is multiplied by a factor of 10 (labeled as MDGL_fixWab10x). This suggests that different tasks exhibit different degree of tolerance to deviations in *w*_*αβ*_.

**Figure S10:**
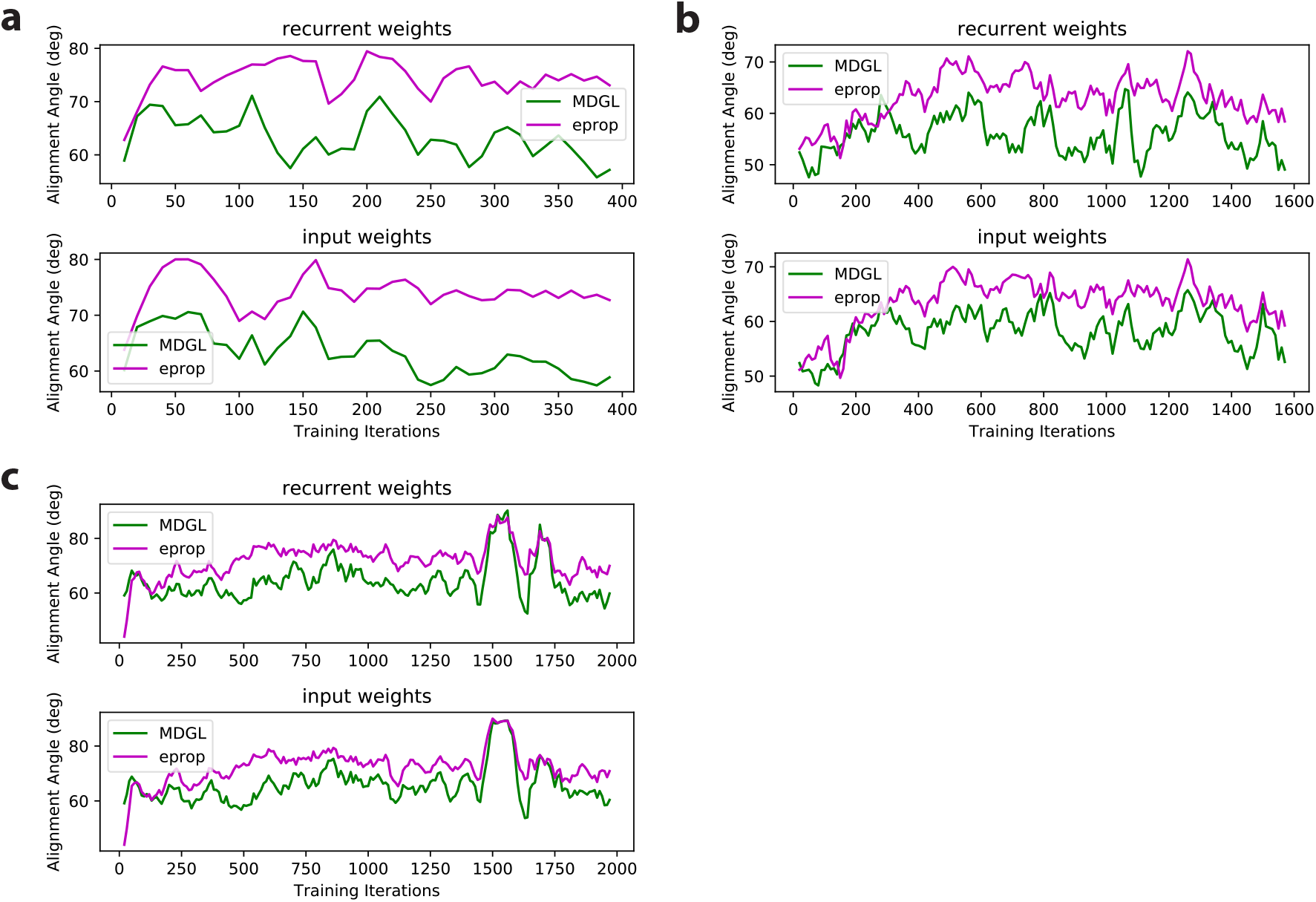
Alignment angle comparison shows that gradients approximated by MDGL is more similar (than e-prop) to the exact gradients. We quantify the similarity between approximated and exact gradients at each time point via the alignment angle, which describes the similarity in the direction of the two update vectors (Supplementary Note 3). This comparison is done for a) pattern generation, b) delayed match-to-sample and c) evidence accumulation tasks. All tasks were trained using MDGL. In each subfigure, alignment angles for recurrent (resp. input) weights over training iterations are shown in the top (resp. bottom) panel.

## Supplementary Tables

**Table S1:** Local modulatory terms of MDGL and TRTRL are significantly aligned

| Task | Beginning |  | Middle |  | End of Training |  |
| --- | --- | --- | --- | --- | --- | --- |
|  | Alignment (°) | Z-score | Alignment (°) | Z-score | Alignment (°) | Z-score |
| Pattern Generation | 36 | -387 | 36 | -385 | 44 | -317 |
| Delayed Match to Sample | 22 | -114 | 48 | -74 | 32 | -99 |
| Evidence Accumulation | 50 | -67 | 44 | -81 | 55 | -59 |

The alignment angles are computed between the local modulatory terms due to neuron-specific rather than type-specific gain (the vectorization of 2D matrices $\widehat{\Gamma}_{pq} = \sum_t \widehat{\Gamma}_{pq,t}$ of TRTRL in Eq. 3 and those due to the cell-type-specific approximation $\Gamma_{pq} = \sum_t \Gamma_{pq,t}$ of MDGL in Eq. 5 using E and I cell types. The angles listed here are for recurrent weight update, and similar values are observed for input weight updates. Here, beginning, middle and end of training refer to after 10, 100 and 500 training iterations, respectively, for the pattern generation task, and after 100, 500 and 2000 training iterations, respectively, for the delayed match to sample and evidence accumulation tasks. Z-scores were computed to show the significance of the difference of alignment from 90°; these scores suggest that TRTRL and MDGL are significantly aligned. Details for computing Z-scores and alignment angles are provided in Supplementary Note 3.

**Table S2:** Local modulatory term of MDGL is not significantly aligned with the approximated gradient by e-prop

| Task | Beginning |  | Middle |  | End of Training |  |
| --- | --- | --- | --- | --- | --- | --- |
|  | Alignment (°) | Z-score | Alignment (°) | Z-score | Alignment (°) | Z-score |
| Pattern Generation | 89 | -8.4 | 80 | -69 | 92 | 16 |
| Delayed Match to Sample | 94 | 5.3 | 88 | -2.3 | 92 | 2.2 |
| Evidence Accumulation | 91 | 1.2 | 91 | 1.4 | 89 | -1.2 |

## Supplementary Note 1 – Online Learning for Leaky Output

Consider a supervised learning task with loss 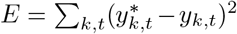 and leaky output 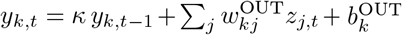, we have the following partial derivative

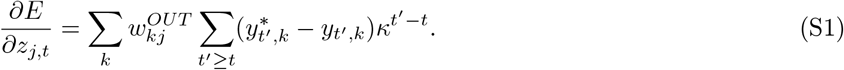

This seemingly provides an obstacle for online learning, because 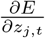 depends on future errors. However, changing the summation order [48]solves this problem and the following derivation can be generalized to the classification task with a simple replacement of 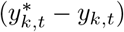:

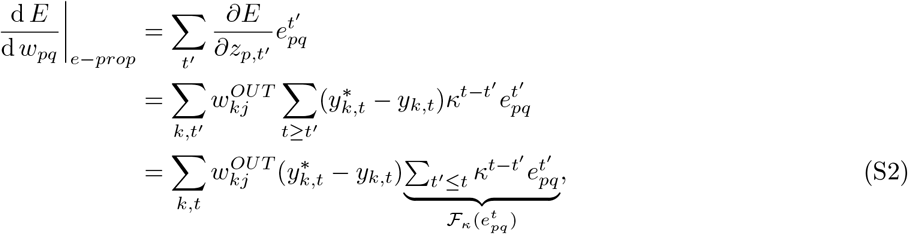

where the order of summations was changed in the last line, and operator *ℱ*_*κ*_ denotes low-pass filtering with ℱ_*κ*_(*x*_*t*_) = *κℱ*_*κ*_(*x*_*t−*1_) + *x*_*t*_. In our actual implementation, we used an exponential smoothing with *ℱ* _*κ*_(*x*_*t*_) = *κℱ*_*κ*_(*x*_*t−*1_) + (1 *− κ*) *∗ x*_*t*_, but dropped the factor (1 *− κ*) in writing for readability.

We also apply the change of summation order trick to the cell-type-specific modulatory signal Γ_*pq,t*_ and assume that activity of neuron *j* is not correlated with the eligibility trace of synapse *pq*:

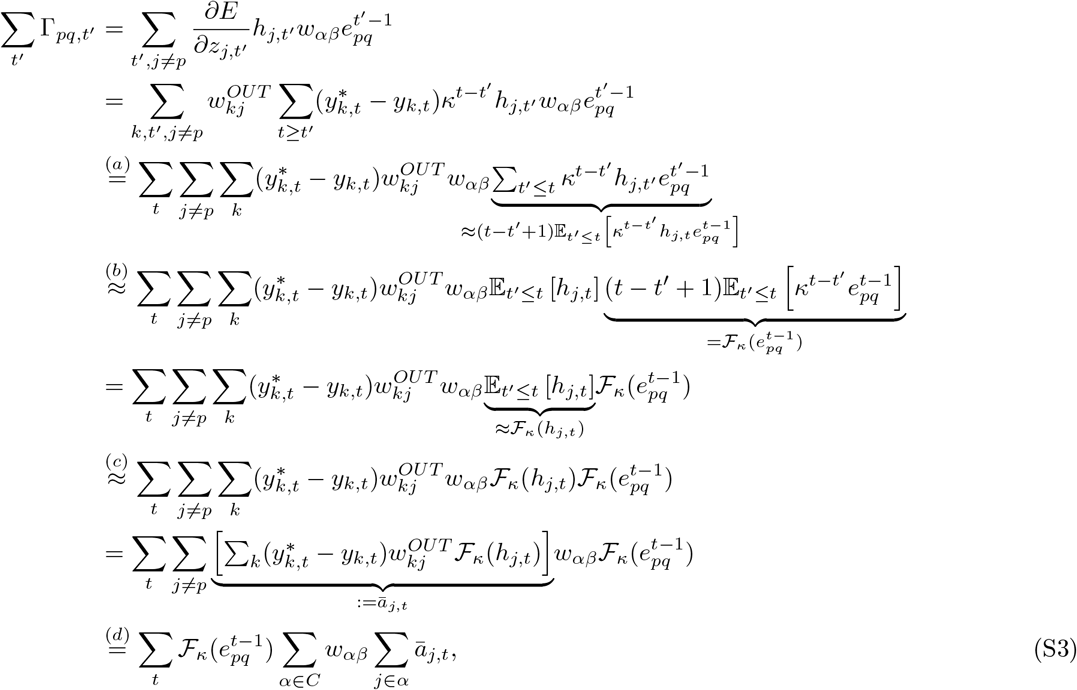

where (*a*) changes the summation order; (*b*) assumes uncorrelatedness between activity *h*_*j,t*_ and 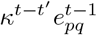 such that 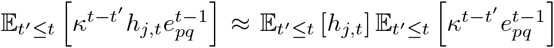; (*c*) approximates the temporal average of *h*_*j,t*_ using an exponential filter 𝔼_*t*_*’≤t* [*h*_*j,t*_] ≈ ℱ_*κ*_(*h*_*j,t*_); (*d*) is a simple change of summation order. We test the validity of above approximation in Figure S3and observe no significant performance degradation due to this approximation.

## Supplementary Note 2 – Detailed Breakdown of MDGL’s Components

In the main text, we stated that our MDGL learning rule combines the eligibility trace with both top-down learning signals and cell-type-specific weighted summation of secreted, diffuse modulators. We so far only expressed these components as derivatives. With the derivation of the online implementation for MDGL in Eq. S3,we are now ready to provide the detailed expressions for each of these components. Combining Eq. S3with Eq. 5and rearranging the summation order gives the following component breakdown for our online approximation to MDGL:

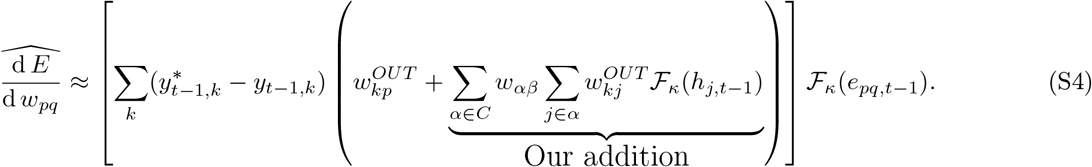

The non-neuron-specific error signal 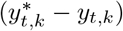 is passed to cells through neuron-specific feedback weights 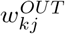, thereby forming neuron-specific learning signal at the receiving end 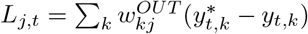. Thus, loosening this neuron-specificity of learning signal can be achieved through approximations to the feedback weights, such as replacing them with random weights [64]or cell-type-specific gains as in Eq. 4. Upon receipt, neuron *j* multiplies *L*_*j,t*_ with ℱ_*κ*_(*h*_*j,t*_), its low-pass filtered activity, and sends the packaged signal *a*_*j,t*_ = *L*_*j,t*_ *ℱ*_*κ*_(*h*_*j,t*_). In updating *w*_*pq*_, our addition allows postsynaptic cell *p* to collect information regarding the activities and learning signals of other cells through cell-type-specific gain *w*_*αβ*_, and combine the received modulatory input with its low-pass filtered eligibility trace.

### Supplementary Note 3 – Analysis and Simulation Details

Throughout this study, the alignment angle *θ* between two vectors, a and b, was computed by *θ* = *acos* (| |*a*^*T*^ *b*| | / | |*a*| | | | *b*| |). The alignment between two 2D matrices was computed by flattening the matrices into vectors. To obtain the significance of alignment in Tables S1and S2,we randomly shuffled the matrices, calculated the resulting alignment angle and repeated for 1000 times to obtain an empirical distribution of alignment angles. The mean *µ* and standard deviation *σ* were computed from the distribution to report the 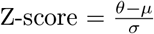.

For spectral analysis, we first performed root mean square normalization on the signal and then computed the power spectral density using Welch’s method [99]. We then found the 3dB frequency by identifying the maximum frequency at which the power is halved from the peak power. For the pattern generation task in Figure 4, our network consisted of 400 LIF neurons. All neurons had a membrane time constant of *τ*_*m*_ = 30ms, a baseline threshold of *v*_th_ = 0.01 and a refractory period of 2ms. Input to this network was provided by 100 Poisson spiking neurons with a rate of 10Hz. The fixed target signal had a duration of 2000ms and given by the sum of five sinusoids, with fixed frequencies of 0.5Hz, 1Hz, 2Hz, 3Hz and 4Hz. For learning, we used mean squared loss function and for visualization, we used normalized mean squared error 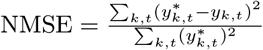 for zero-mean target output 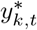. All weight updates were implemented using Adam with default parameters [100]and a learning rate of 1 *×* 10^*−*3^. In addition, we applied firing rate regularization with *c*_reg_ = 10 and *f* ^target^ = 10*Hz*.

For the delayed match to sample task in Figure 5, our network consisted of 50 LIF neurons and 50 ALIF neurons (100 LIF neurons and 80 ALIF neurons for the alternative setup in Figure S5). All neurons had a membrane time constant of *τ*_*m*_ = 20ms, a baseline threshold of *v*_th_ = 0.01 and a refractory period of 5ms. The time constant of threshold adaptation was set to *τ*_*b*_ = 1400ms, and its impact on the threshold was set to *β* = 1.8. Input to this network was provided by three populations, as illustrated in Figure 5B. The first (resp. second) population consisted of 20 units and produced Poisson spike trains with a rate of 40Hz when the first (resp. second) cue takes a value of 1, otherwise it stays quiescent. The last input population of 10 units produced Poisson spike trans of 10Hz throughout the trial in order to prevent the network from being quiescent during the delay. For the alternative setup in Figure S5,the first (resp. second) input population produced Poisson spike trains with a rate of 40Hz when cue 1 (resp. cue 2) is presented, otherwise it fires at 10Hz. For learning, we used cross-entropy loss function and the target corresponding to the correct output was given at the end of the trial. As done in the evidence accumulation task, a weight update was applied once every 64 trials and the gradients were accumulated during those trials additively. All weight updates were implemented using Adam with default parameters [100]and a learning rate of 2.5 *×* 10^*−*3^. In addition, we applied firing rate regularization with *c*_reg_ = 0.1 and *f* ^target^ = 10Hz.

For the evidence accumulation task in Figure 6, our network consisted of 50 LIF neurons and 50 ALIF neurons. All neurons had a membrane time constant of *τ*_*m*_ = 20ms, a baseline threshold of *v*_th_ = 0.01 and a refractory period of 5ms. The time constant of threshold adaptation was set to *τ*_*b*_ = 2000ms, and its impact on the threshold was set to *β* = 1.8. Input to this network was provided by four populations of 10 neurons each, as illustrated in Figure 6B. The first (resp. the second) population produced Poisson spike trains with a rate of 40Hz when a cue was presented on the left (resp. right) side of the track. The third input population spiked randomly through the decision period with a firing rate of 40Hz and was silent otherwise. The last input population produced Poisson spike trains with a rate of 10Hz throughout the trial in order to prevent the network from being quiescent during the delay. For learning, we used the cross-entropy loss function and the target corresponding to the correct output was given at the end of the trial. A weight update was applied once every 64 trials and the gradients were accumulated during those trials additively. All weight updates were implemented using Adam with default parameters [100]and a learning rate of 2.5 *×* 10^*−*3^. In addition, we applied firing rate regularization with *c*_reg_ = 0.1 and *f* ^target^ = 10*Hz*. For all simulations, we used a time step of 1ms, as done in [48]. We also assumed a synaptic delay of 1ms for all synapses.

## Notes

### Competing Interest Statement

The authors have declared no competing interest.

### Summary of Updates

- Modified introduction - Additional supplementary figures

